# Glioma-induced DNMT3A-dependent reduction of DNA methylation in microglia promotes a transient anti-tumoral phenotype

**DOI:** 10.1101/2025.02.22.639606

**Authors:** Mathilde Cheray, Adriana-Natalia Murgoci, Adamantia Fragkopoulou, Carlos F.D. Rodrigues, Martin Škandík, Ahmed M. Osman, Christine Hong, Guillermo Vazquez-Cabrera, Lara Friess, Lena-Maria Carlson, Shigeaki Kanatani, Yue Li, Anastasius Damdimopoulos, Per Uhlén, Fredrik Kamme, Klas Blomgren, Bertrand Joseph

**Author notes:** Correspondence should be addressed to B.J. These authors contributed equally (Adriana-Natalia Murgoci, Adamantia Fragkopoulou, Carlos F.D. Rodrigues).

## Abstract

Glioblastoma, aggressive primary brain tumors with a dismal prognosis, promote the recruitment of microglia, brain resident innate immune cells, and ultimately their activation toward a tumor-supportive phenotype that increases gliomal proliferation and invasion capability. Here, we report that upon stimulation by glioma cells, microglia transit via a reactive state holding anti-tumoral properties coupled to reduced DNMT3A chromatin occupancy and DNA demethylation that promote microglial pro-inflammatory gene expressions. We find that upon repression of *Dnmt3a* expression in microglia, those cells maintain anti-tumoral attributes *in vitro* and *in vivo*. In a syngeneic immunocompetent glioblastoma mouse model, brain delivery of antisense oligonucleotide targeting *Dnmt3a* expression led to reduced tumor growth. Taken together, our results reveal the involvement of DNA demethylation in the control of glioma cells-induced microglia activation and indicate that microglial DNMT3A is a potentially therapeutic target to treat brain neoplasms such as glioblastoma that include a microglial component.

## Introduction

Glioblastoma, IDH-wildtype (GB), are highly aggressive primary brain tumors with limited therapeutic options, and a dismal prognosis for patients.^1^ The intrinsic capacity of these high-grade gliomas to infiltrate the surrounding brain tissue impedes surgical resection, and contributes significantly to the failure of current therapeutic treatments, predictably results in high rates of early recurrence. Despite multimodal therapy with the concomitant use of radiation and adjuvant treatment (*e.g.* temozolomide, bevacizumab), the overall median survival for patients with GB is still restricted to about 15 months.^1^ Thus, primary high-grade gliomas, such as GB, with their significant mortality and morbidity remain one of the major challenges of today’s oncology.

One emerging view is that if new therapeutic strategies are to be found for this devastating neoplasm, one will have to take into account that GB tumors are heterogeneous with respect to the composition of bona fide tumors cells and with respect to a range of intermingling non-neoplastic cells which also play a vital role in controlling the course of the disease.^2–4^ In fact, GB tumors contain an important infiltration of immune cells, especially myeloid cells are enriched.^4^ This robust myeloid-dependent immune response in GB is thought to increase tumor aggressiveness and lead to reduced survival.^5^ The tumor-associated myeloid cells are composed of microglia, *i.e*., resident immune cells of the central nervous system, and of peripherally recruited bone marrow-derived macrophages (BMDM). The locations of these two myeloid cell types differ, at least in an established human GB tumor, arguing for distinct functions: BMDM are found in the core of the tumor, whereas microglia are principally found in its periphery.^6–9^ In mouse models of the disease, BMDM were shown, following disruption of the blood-brain barrier^10^, to infiltrate these tumors only at later stages of their expansion.^3, 11^ In fact, compelling studies in GB mouse models that used head-protected irradiation chimeras, as well as multiple genetic lineage tracing systems, demonstrated that resident microglia represent the predominant and early source of myeloid cells within GB. ^3, 11, 12^ As further evidence of their essential role in glioma progression, removal of microglia, both in brain organotypic slices and genetic mouse models, inhibited glioma invasiveness and even impacted on tumor angiogenesis.^2,13, 14^

Targeting the resident microglia, as well as the peripherally recruited BMDM, in the tumor microenvironment has been proposed as an intervention to combat GB expansion. Myeloid cells depend on colony stimulating factor-1 (CSF1) for differentiation and survival. Blockade of the CSF1 receptor (CSF1R) initially yielded promising results in a mouse model of GB with increased survival and regression of tumors.^15^ However, GB tumors are shown to reoccur due the acquisition by the tumor associated microglia/macrophages of a resistance to CSF1R inhibition and the production of insulin growth factor 1 (IGF1) that instead drive tumor growth and survival. This could represent a mechanism that could account for the disappointing results observed in clinical trials of CSF1R blockade.^16, 17^

During the course of the disease, microglia are recruited by the glioma cells as immune competent cells that potentially exhibit anti-tumoral effects, before they are reprogrammed into tumor-supporting cells by the same cancer cells, hence defining different microglial reactive states with unique targetable functions.^2, 18, 19^ Therefore, an additional strategy for targeting microglia in the tumor microenvironment revolve around the reprogramming of these cells. This approach relies on either impairing the acquisition of tumor-promoting capacity by microglia or on converting them into tumor-attacking cells, maintaining some of their intrinsic properties as innate immune cells. However, the molecular mechanisms that control the transition of microglia from an immune reactive state with a unique gene expression profile and possible anti-tumoral properties into a phenotype with a distinct transcriptome that is associated with tumor-trophic functions in response to cues from glioma cells, remain elusive. Transcriptional activation and repression are strongly influenced by and associated with changes in chromatin structure. In fact, epigenetic changes, including histone posttranslational modifications and DNA methylation, are important regulators of gene expression, and have been involved in cell phenotype regulation as well as cellular reprogramming.^20–22^ DNA methylation patterns placed by the DNA methyltransferase (DNMT) enzymes are reversible heritable marks conserved during cell division, which have been shown to be actively involved in cell reprogramming processes.^21, 22^ We sought to determine whether DNMT-mediated DNA methylation could be involved in glioma-induced microglial activation. We found that in response to glioma stimuli, a DNMT3A-dependent transient downregulation of microglial DNA methylation contributed to the acquisition of a unique transcriptome associated with a microglial reactive state with anti-tumoral properties. Repression of microglial *Dnmt3a* expression *in vivo* enabled the activation of microglia cells and led to a reduction of GB tumor growth in a syngeneic mouse model.

## Results

### Microglia are initially activated as immune reactive cells in response to glioma stimulation

Glioma cells are reported to reprogram microglia, the brain resident myeloid immune cells, towards a tumor-supporting phenotype (reviewed in ^2, 23, 24^). As an illustration and as previously reported^19, 25^, in a transwell migration assay, BV2 microglia stimulated by soluble factors originating from C6 glioma cells, are able to promote the invasion capabilities of the cancer cells assayed at 24 hours (Fig.1a). We hypothesized that in response to stimulation by glioma cells, microglia could transit through distinctive states of activation(s) before acquiring a tumor-supportive phenotype.

**Figure 1.**
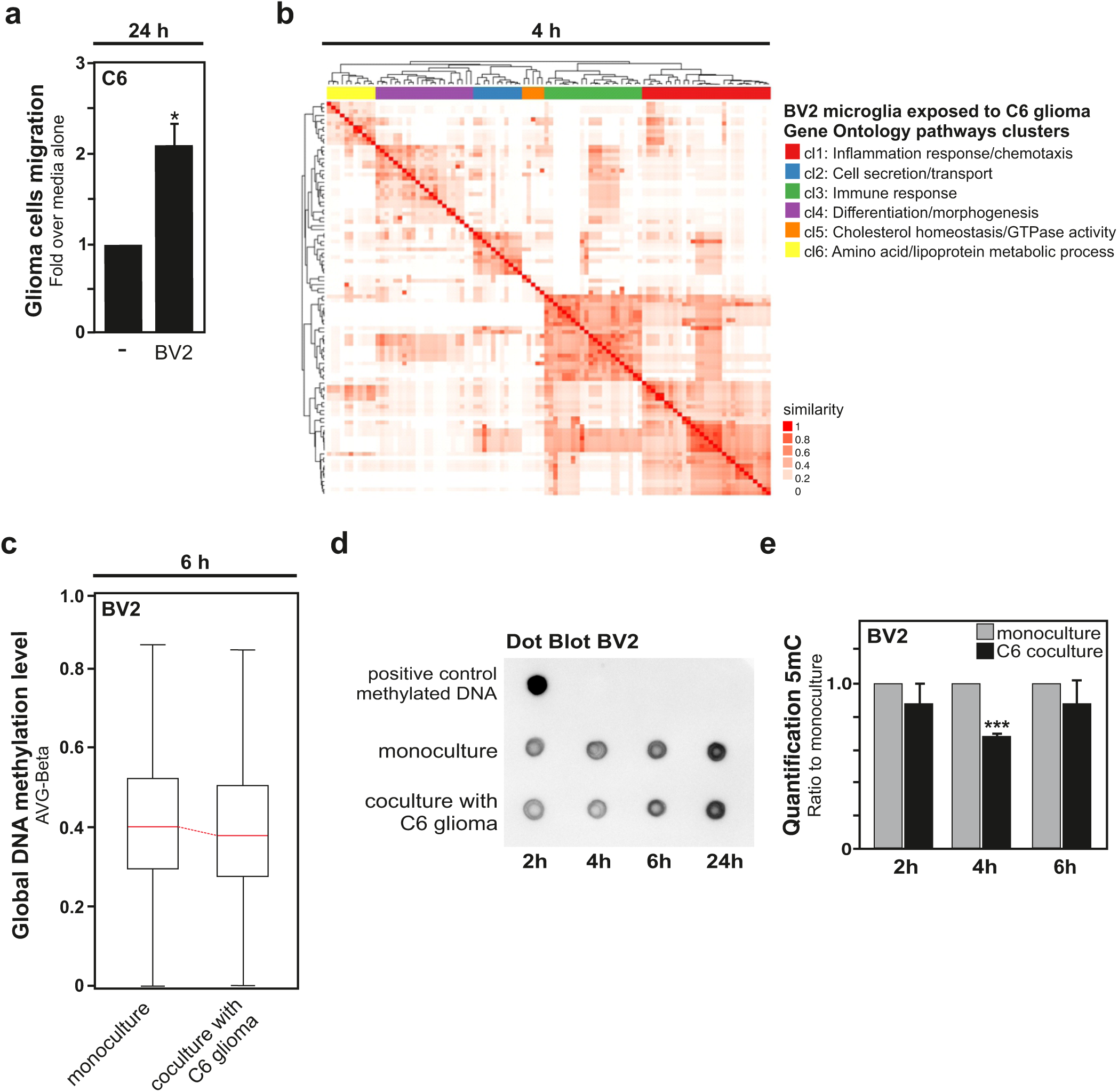
Short term coculture of microglia with glioma cells induces an immune response associated transcriptome and a reduction of global DNA methylation in microglia. **a**, BV2 microglia improve the migration capability of C6 glioma cells tested at 24 hour using a transwell migration assay. For this assay, C6 glioma cells seeded in inserts in 5% medium were placed in plates containing 10% FBS medium (-control condition) or BV2 microglia in 10% FBS medium (+ BV2 condition). **b**, RNA-seq clustering analysis of the top 100 most significant Gene Ontology (GO) pathways found upregulated in BV2 microglia cocultured with C6 glioma cells for 4 hours compared to BV2 microglia in monoculture using the weight01 algorithm. The intensity of the red color shows the degree of similarity and the dendrograms illustrate the hierarchical clustering calculation based on the distance or similarity between rows. The majority of the GOs are associated with the immune system and form one cluster related to “inflammatory response/chemotaxis” (cluster 1, colored in red) and one linked to “immune response” (cluster 3, colored in green). **c**, High-throughput profiling of DNA methylation status at CpG islands performed in BV2 microglia after 6 hours monoculture *versus* 6 hours coculture with C6 glioma cells. The average methylation levels (Beta values) on the CpG islands for each sample obtained from the Illumina^®^ Infinium HumanMethylation450 BeadChip array analysis are shown as a box plot. **d**,**e**, Dot Blot analysis of 5-methylcytosine content (5-mC) in BV2 microglia after different time points in monoculture or coculture with C6 glioma cells (**d**) with quantification (**e**), presented as fold of monoculture condition for each time point set as 1 (n=4; Data are shown as mean ± sem).

To investigate this possibility, BV2 microglia were exposed for limited time, *i.e*., 2 or 4 hours, to soluble factors originating from C6 glioma cells in a segregated coculture set-up, thereafter microglia were collected, and bulk RNA sequencing (RNA-seq) was performed. The genome wide analysis of the microglial transcriptome showed differentially expressed genes (DEGs) between microglia collected from monoculture, used as control, and those collected from segregated coculture with glioma cells. Significantly more DEGs (p-value <0.05) were found in microglia after 4 hours (199 DEGs) compared to 2 hours (32 DEGs) coculture with glioma cells. Most of the microglial genes found to be regulated at the 2 hours’ time-point were also found to be affected at the 4 hours’ time-point (supplementary data file). Volcano plots of DEGs between unstimulated microglia (monoculture) and glioma-stimulated microglia (coculture for 2 or 4 hours) are depicted in Extended Data Fig.1a,b.

When the genes found to be upregulated in microglia after 4 hours exposure to glioma cells were analyzed for Gene Ontology (GO) pathways clustering, one cluster related to “inflammatory response” (cluster 1, colored in red) and one linked to “immune response” (cluster 3, colored in green) were found to be significantly enriched. These two early activated gene clusters are associated with the immune system, along with a microglial pro-inflammatory phenotype (Fig.1b). Further analysis of the first 50 significantly enriched GO pathways in microglia-glioma coculture vs microglia monoculture condition, showed that 20 of these GO pathways are in fact associated with an immune response of the microglial cells (Table 1). In addition, Cytoscape GO network analyses performed on genes found to be upregulated in microglia after 2 or 4 hours exposure to glioma cells further supported the concept of an immune reactive microglial phenotype at those early stages of glioma-induced microglia activation with an observed enrichment for clusters linked to biological processes associated with the regulation of immune- and stress responses (Extended Data Fig.1c,d).

**Table 1.**
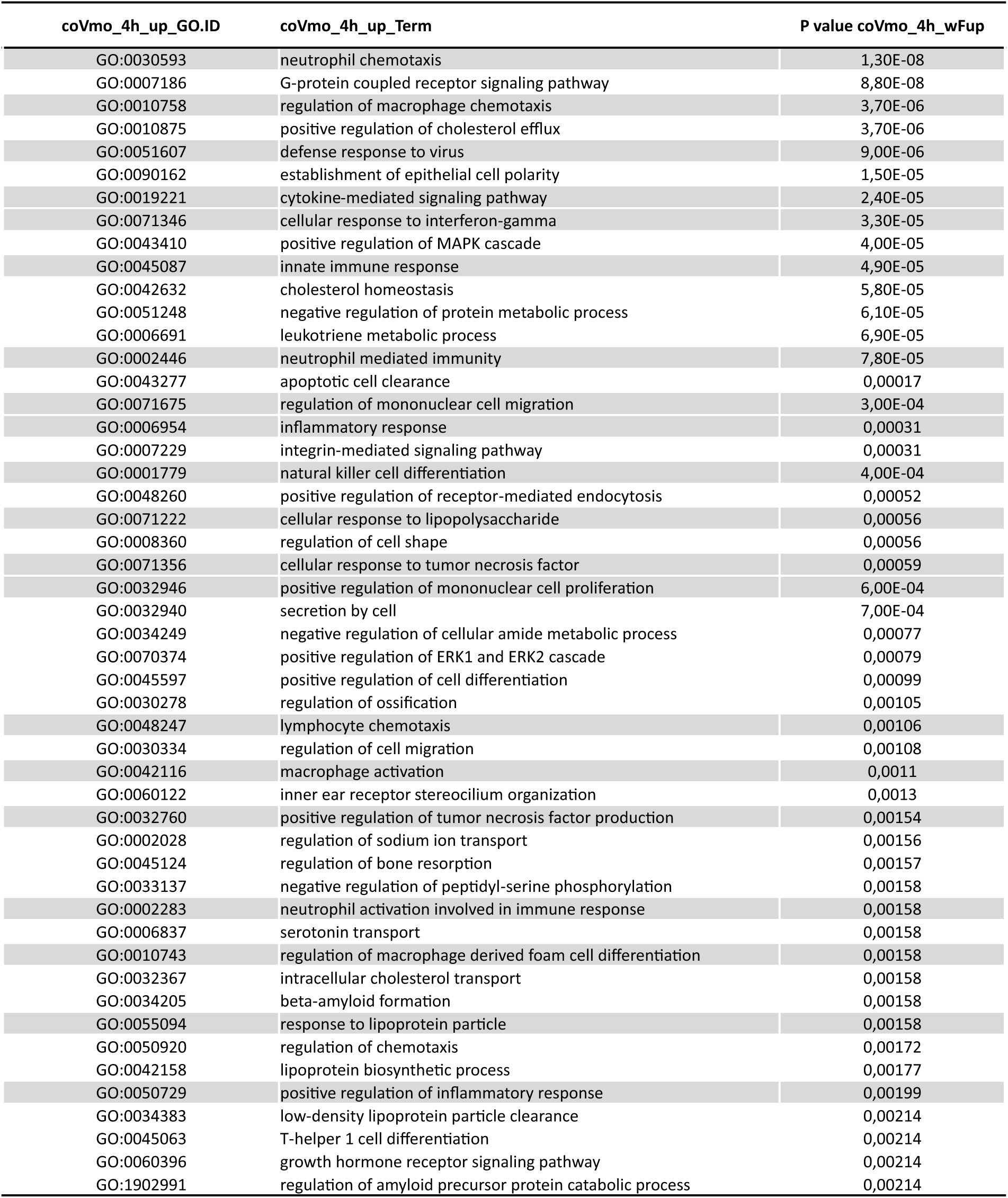
Top 50 Gene Ontology pathways (GO) overexpressed in BV2 cells in coculture with C6 at 4h compared to monoculture condition (The GO pathways highlighted in grey are related to pro-inflammatory cell phenotype).

| coVmo_4h_up_GO.ID | coVmo_4h_up_Term | P value coVmo_4h_wFup |
| --- | --- | --- |
| GO:0030593 | neutrophil chemotaxis | 1,30E-08 |
| GO:0007186 | G-protein coupled receptor signaling pathway | 8,80E-08 |
| GO:0010758 | regulation of macrophage chemotaxis | 3,70E-06 |
| GO:0010875 | positive regulation of cholesterol efflux | 3,70E-06 |
| GO:0051607 | defense response to virus | 9,00E-06 |
| GO:0090162 | establishment of epithelial cell polarity | 1,50E-05 |
| GO:0019221 | cytokine-mediated signaling pathway | 2,40E-05 |
| GO:0071346 | cellular response to interferon-gamma | 3,30E-05 |
| GO:0043410 | positive regulation of MAPK cascade | 4,00E-05 |
| GO:0045087 | innate immune response | 4,90E-05 |
| GO:0042632 | cholesterol homeostasis | 5,80E-05 |
| GO:0051248 | negative regulation of protein metabolic process | 6,10E-05 |
| GO:0006691 | leukotriene metabolic process | 6,90E-05 |
| GO:0002446 | neutrophil mediated immunity | 7,80E-05 |
| GO:0043277 | apoptotic cell clearance | 0,00017 |
| GO:0071675 | regulation of mononuclear cell migration | 3,00E-04 |
| GO:0006954 | inflammatory response | 0,00031 |
| GO:0007229 | integrin-mediated signaling pathway | 0,00031 |
| GO:0001779 | natural killer cell differentiation | 4,00E-04 |
| GO:0048260 | positive regulation of receptor-mediated endocytosis | 0,00052 |
| GO:0071222 | cellular response to lipopolysaccharide | 0,00056 |
| GO:0008360 | regulation of cell shape | 0,00056 |
| GO:0071356 | cellular response to tumor necrosis factor | 0,00059 |
| GO:0032946 | positive regulation of mononuclear cell proliferation | 6,00E-04 |
| GO:0032940 | secretion by cell | 7,00E-04 |
| GO:0034249 | negative regulation of cellular amide metabolic process | 0,00077 |
| GO:0070374 | positive regulation of ERK1 and ERK2 cascade | 0,00079 |
| GO:0045597 | positive regulation of cell differentiation | 0,00099 |
| GO:0030278 | regulation of ossification | 0,00105 |
| GO:0048247 | lymphocyte chemotaxis | 0,00106 |
| GO:0030334 | regulation of cell migration | 0,00108 |
| GO:0042116 | macrophage activation | 0,0011 |
| GO:0060122 | inner ear receptor stereocilium organization | 0,0013 |
| GO:0032760 | positive regulation of tumor necrosis factor production | 0,00154 |
| GO:0002028 | regulation of sodium ion transport | 0,00156 |
| GO:0045124 | regulation of bone resorption | 0,00157 |
| GO:0033137 | negative regulation of peptidyl-serine phosphorylation | 0,00158 |
| GO:0002283 | neutrophil activation involved in immune response | 0,00158 |
| GO:0006837 | serotonin transport | 0,00158 |
| GO:0010743 | regulation of macrophage derived foam cell differentiation | 0,00158 |
| GO:0032367 | intracellular cholesterol transport | 0,00158 |
| GO:0034205 | beta-amyloid formation | 0,00158 |
| GO:0055094 | response to lipoprotein particle | 0,00158 |
| GO:0050920 | regulation of chemotaxis | 0,00172 |
| GO:0042158 | lipoprotein biosynthetic process | 0,00177 |
| GO:0050729 | positive regulation of inflammatory response | 0,00199 |
| GO:0034383 | low-density lipoprotein particle clearance | 0,00214 |
| GO:0045063 | T-helper 1 cell differentiation | 0,00214 |
| GO:0060396 | growth hormone receptor signaling pathway | 0,00214 |
| GO:1902991 | regulation of amyloid precursor protein catabolic process | 0,00214 |

### Early glioma cells-induced microglia activation is associated with a transient reduction in global microglial DNA methylation

Methylation of the 5′-position of cytosine residues, located adjacent to a guanosine (CpG), is a reversible covalent modification of DNA resulting in the production of 5-methyl-cytosine (5-mC) that changes the biophysical characteristic of the DNA. In mammals, DNA methylation, is recognized as an epigenetic process, known to efficiently influence gene expression at the transcriptional level.^26^ Whereas the contribution of changes in histone posttranslational modifications, another epigenetic mechanism regulating transcriptional responses, to the polarization of microglia toward a tumor trophic phenotype by glioma cells has been looked,^20, 25, 27, 28^ the possible contribution of DNA methylation remained to be fully explored. Hence, we speculated that the rapid microglial transcriptomic immune reactive-related response observed could be due to changes in DNA methylation. Using the above-described segregated coculture set-up, DNA methylation level was examined in BV2 microglia stimulated with soluble factors originating from C6 glioma cells. We took advantage of the Illumina^®^ Infinium Methylation 450 BeadChip array which gives the methylation status of over 450,000 methylation sites per sample at single nucleotide resolution. Remarkably, global DNA methylation level was found to be reduced in the microglia after 6 hours of segregated coculture with the glioma cells (Fig.1c). DNA dot blot analysis of 5-mC content in BV2 microglia exposed to C6 glioma cells further revealed that the reduction in global DNA methylation in these cells was transient. Indeed, while significant reduction in microglial global DNA methylation level was observed after 4 hours of segregated coculture with glioma cells, the 5-mC levels were found to rise back to the level observed in microglial monoculture at 24 hours (Fig.1d, e).

### Early response of microglia to glioma cells leads to the expression of genes associated with a pro-inflammatory and motile cell phenotype

To further explore the prospect that the microglial reactive state, and thereby potential associated functions, could change over time upon activation by glioma cells, the expression of genes which are commonly used for the characterization of different microglial phenotypes was investigated by RTqPCR in a time-dependent manner (upon exposure to glioma cells for 2, 4 and 6 hours). Early and sustained mRNA expression for two markers associated with the microglial tumor-supportive phenotype, *i.e.* chemokine (C-C Motif) ligand 22 (*Ccl22*), and chitinase-like protein 3 (*Chil3*, also known as *Ym1*), was observed in BV2 microglia upon coculture with C6 glioma cells^19^ (Fig.2a). Early induction of matrix metalloproteinase-14 (membrane-inserted) (*Mmp14*, also known as *Mt1-mmp*) microglial mRNA expression, but not *Mmp9*, was also observed in microglia, but its expression appears to be transient (Fig.2b). The expression of MMP14 in microglia has been associated with the recruitment of microglia into the glioma tumor.^13, 29^ Expression of genes encoding for the cytokines interleukin 1β (*Il1β*) and interleukin 6 (*Il6*) was also examined in C6 glioma cells stimulated BV2 microglia (Fig.2c). IL1β is a prototypic pro-inflammatory cytokine that plays key roles in both acute and chronic inflammatory responses.^30^ *Il6* gene expression is relevant in a clinical context, as elevated IL6 expression is associated with poor survival of patients with glioma.^31^ IL6 is an interleukin reported to acts as both a pro- and anti-inflammatory cytokine. IL6 signaling seems to contribute to glioma malignancy through the promotion of glioma stem cell growth and survival.^31^ Moreover, IL6 participates in the maintenance of the microglial tumor-supportive functions.^32^ Early induction of microglial mRNA expression for both cytokines was observed (Fig.2c). *Il6* mRNA expression levels were still found to be significantly upregulated at the 6 hours’ time-point. In contrast, the messenger expression for the pro-inflammatory cytokine *IL1β*, was shown to be limited in time and could only be observed in BV2 microglia at the 2 hours’ time-point upon coculture with C6 glioma cells (Fig.2c). Early induction of *IL1β*, but not *Il6* gene expression, was also observed in the human microglial cell line HMC3 upon segregated coculture with the human U87-MG glioma cells (Extended Data Fig.2a).

**Figure 2.**
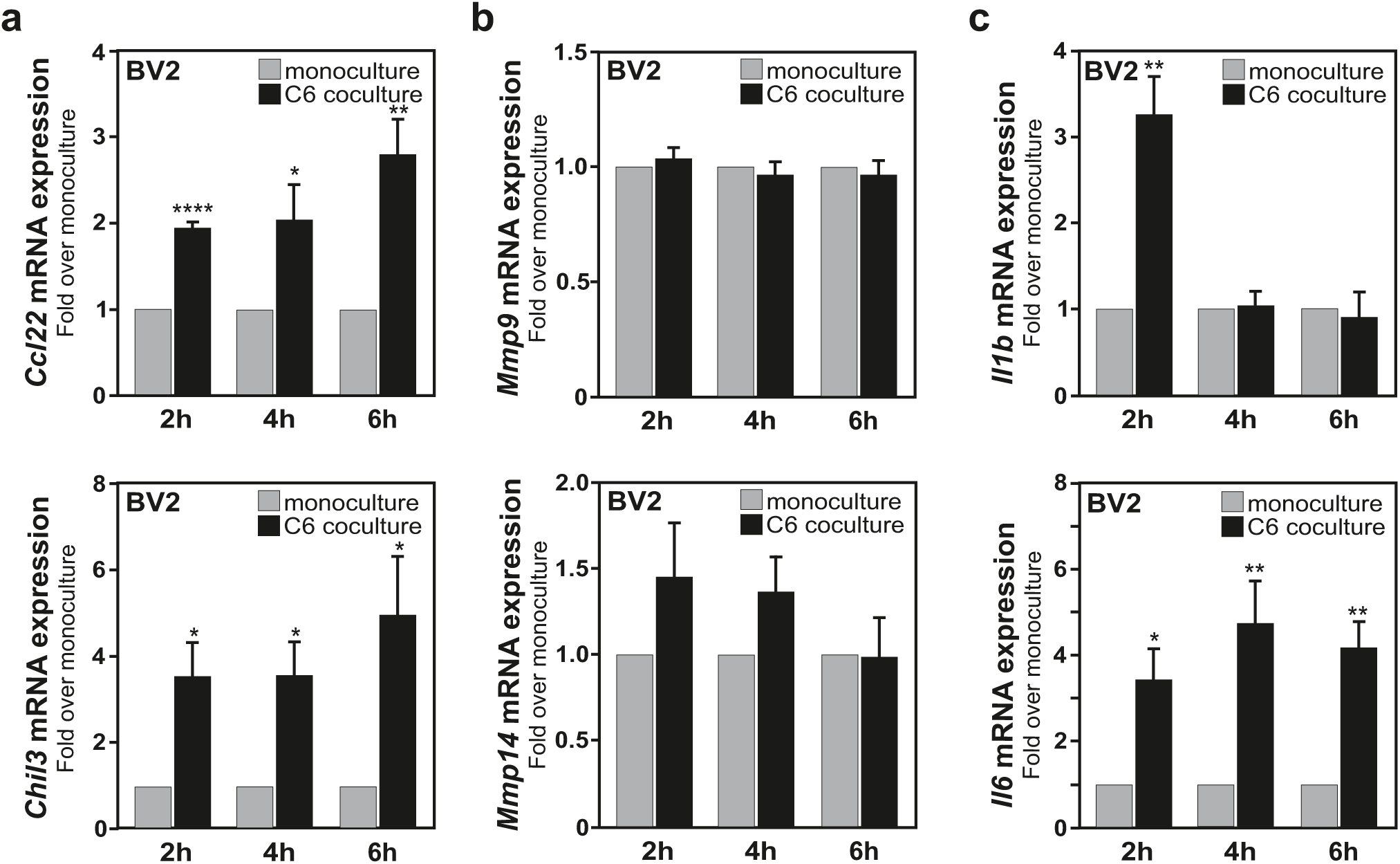
Short term coculture of microglia with glioma cells induces the microglial expression of genes associated to both a tumor-supportive phenotype and a proinflammatory phenotype. **a-c**, RTqPCR analysis of mRNA expression levels in BV2 microglia exposed to coculture with C6 glioma cells for 2, 4, and 6 hours of genes frequently associated to a microglial tumor-supportive phenotype, *i.e*., *Ccl22* and *Chil3/Ym1* (**a**), matrix metalloproteinases, *i.e*., *Mmp9* and *Mmp14/Mt1-mmp* (**b**), and proinflammatory cytokines, *i.e*., *Il1b* and *Il6* (**c**). (n=3; Data are shown as mean ± sem).

Treatment with lipopolysaccharide (LPS), the major component of Gram-negative bacterial walls and a ligand for microglial Toll-like receptor 4 (TLR4), leads to microglial activation toward a pro-inflammatory phenotype with associated neurotoxicity.^33^ Microglia also express IL4 receptors and, when these brain resident myeloid cells are exposed to this interleukin, they acquire instead anti-inflammatory properties.^34^ In addition, different gliomas secrete IL4, and IL4 can induce microglia/macrophages to support tumor survival by creating an immunosuppressive environment.^35^ Treatment with IL4 is reported to promote a microglial phenotype resembling the tumor supportive one. The response of BV2 microglia to LPS and IL4 treatment are characterized respectively by the induction of nitric oxide synthase 2 (NOS2, also known as inducible NOS, iNOS) and arginase 1 (ARG1) protein expressions, respectively, which are otherwise absent in the unstimulated microglia (Extended Data Fig.3a).^19, 33^ Noteworthy, LPS or IL4 treated-BV2 microglia show a different pattern of expression for the *IL1β* gene as compared to microglia responding to a stimulation by glioma cells, confirming a specific regulation of the microglial *IL1β* gene upon exposure to glioma cells (Extended Data Fig.3c,d).

### Decrease in microglial DNA methylation is associated with transient downregulation of DNMT3A and DNMT3B

DNA methylation is catalyzed by DNA methyltransferases (DNMTs), enzymes involved in the covalent transfer of a methyl group to the fifth carbon atom of a cytosine ring.^21, 22^ Therefore, the protein expression level of the main DNMTs, *i.e.* DNMT1, DNMT3A and DNMT3B, was investigated in BV2 microglia upon exposure to C6 glioma cells. Immunoblot analysis uncovered a significant reduction in the expression level of microglial DNMT3A and DNMT3B proteins between 4 and 6 hours segregated coculture with the glioma cells. Likewise, the observed decrease in microglial global DNA methylation, the reduction in DNMT3A and DNMT3B protein expression was transient, and their expression levels at 24 hours in microglia from segregated glioma coculture were undistinguishable from microglia in monoculture (Fig.3a,b). In those conditions, no significant reduction in the DNMT1 protein expression levels was observed (Fig.3a,b). Of interest, a significant increase in microglial *Dnmt3a* and *Dnmt3b* mRNA expression levels was observed after 6 hours of microglia-glioma coculture that could be explained as a compensatory mechanism activated by the microglia to recover from the reduction in protein expression (Fig.3c). Decreased microglial DNMT3A protein and *DNMT3A* mRNA expression upon exposure to glioma derived soluble factors was further confirmed with additional segregated coculture combination with the human HMC3 microglia cell line and the human U87-MG glioma cell line (Extended Data Fig.2b-d). DNMT3A and DNMT3B are highly homologous proteins, and share overlapping functions; however, they also exert distinct functions.^36–38^ Of interest for the current study, a conditional *Dnmt3a* knockout mouse model revealed that large conserved domains of low DNA methylation, which cover regions enriched for genes involved in transcriptional regulation and therefore potentially with cell differentiation, are specifically maintained by DNMT3A.^39^ In human, *DNMT3A* haploinsufficiency causes DNA methylation defects that lead to abnormal gene expression and an impaired immune response of myeloid cells, as well as predisposition to myeloid malignancies.^40, 41^ In mouse expressing a histone H3.3G34R amino acid substitution, *i.e*., a human germline mutation reported to cause severe neurodevelopmental disorders, DNMT3A recruitment to the chromatin is impaired, and an accumulation of disease-associated microglia coupled with the expression of innate immune-related genes is observed.^42^ In fact, *DNMT3A* alterations are frequently associated with human myeloid malignancies (reviewed in ^43^). Collectively, the aforementioned observations and studies make DNMT3A an interesting candidate for the regulation of a specific microglial reactive state induced by early exposure to glioma cells.

**Figure 3.**
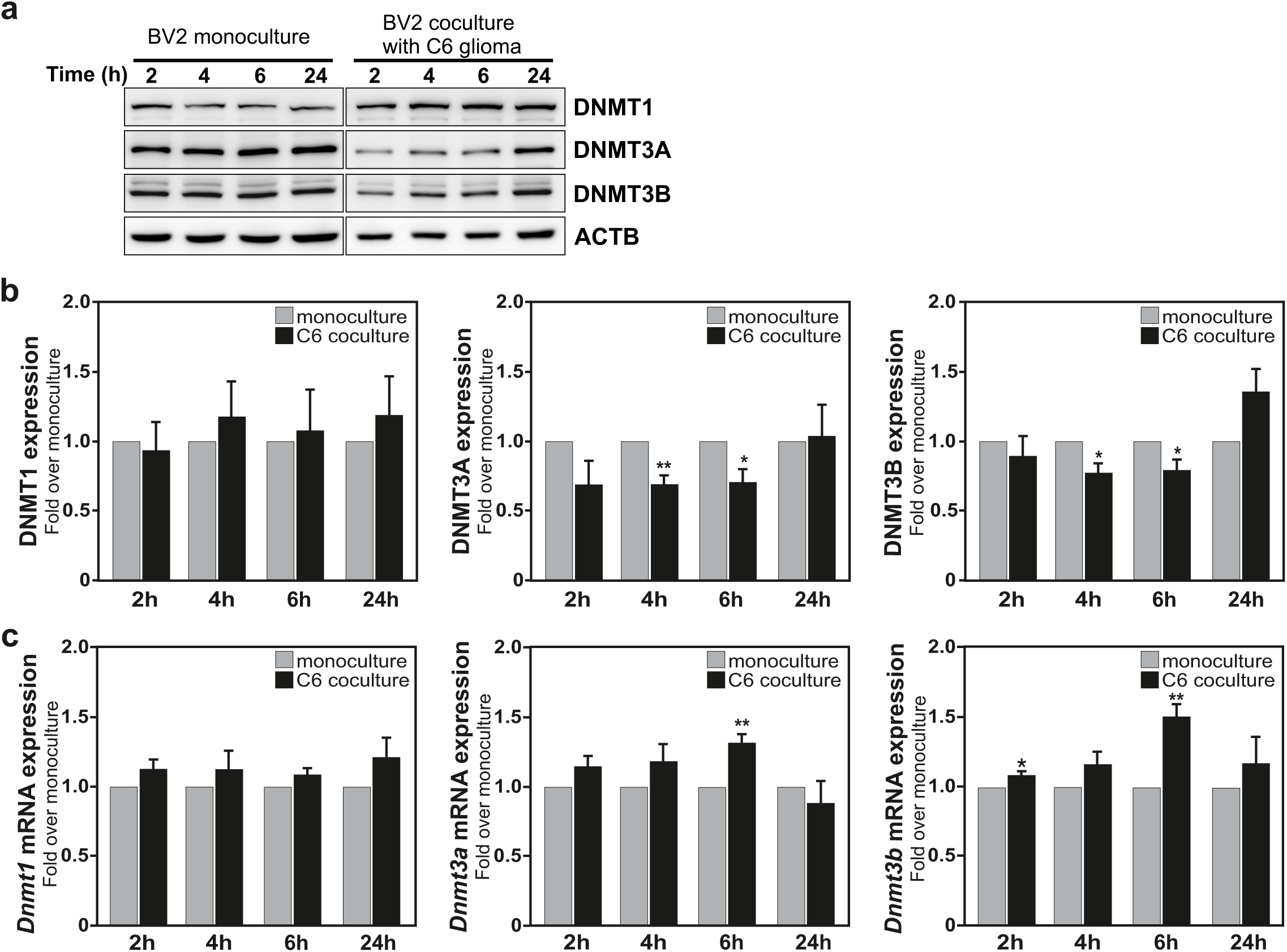
Transient reduction in DNMT3A/3B expression levels observed in BV2 microglia stimulated by C6 glioma cells. **a**,**b**, Immunoblot analysis (**a**), and quantifications (**b**), of DNMTs proteins (DNMT1, DNMT3A and DNMT3B) and β-actin (ACTB) expression levels in BV2 microglia in monoculture or in coculture with C6 glioma cells at the indicated time-points (n=3; Data are shown as mean ± sem). **c**, RTqPCR analysis of microglial *Dnmt1*, *Dnmt3a* and *Dnmt3b* mRNA expression levels, using identical conditions as the ones described in panel a (n=3; Data are shown as mean ± sem).

Thereafter, we wanted to explore whether the observed transient DNMT3A downregulation in microglia cells is a distinctive characteristic of their activation by glioma cells, or a general mark for reactive microglia. To address this question, BV2 microglia were treated with LPS, or IL4. Neither LPS treatment, nor IL4 treatment, had an effect on the DNMT3A protein expression (Extended Data Fig.3a,b). However, these treatments appear to affect microglial *Dnmts* mRNA expression levels (Extended Data Fig.3e,f).

Collectively, these data demonstrate that the transient reduction in DNMT3A expression level and associated decrease in global DNA methylation are distinctive characteristics of the activation of microglia by gliomas cells.

### Transient reduction in DNMT3A DNA occupancy is observed in glioma-stimulated microglia

DNMT3A chromatin immunoprecipitation sequencing (ChIP-seq) was performed to monitor at the genome-wide level the DNA binding of this enzyme upon glioma-induced microglia activation. This analysis revealed a general reduction in DNMT3A DNA occupancy in microglia stimulated for 2 hours by glioma cells. This reduction in DNMT3A DNA binding was sustained at 4 hours while at a lower level (Fig.4a). Remarkably, the combined analysis of the RNA-seq and DNMT3A ChIP-seq microglial data sets showed that reduced DNMT3A DNA occupancy was observed (at 2 hours) in genes whose expressions were found to be upregulated (at 4 hours) upon stimulation by glioma cells (Fig.4b).

**Figure 4.**
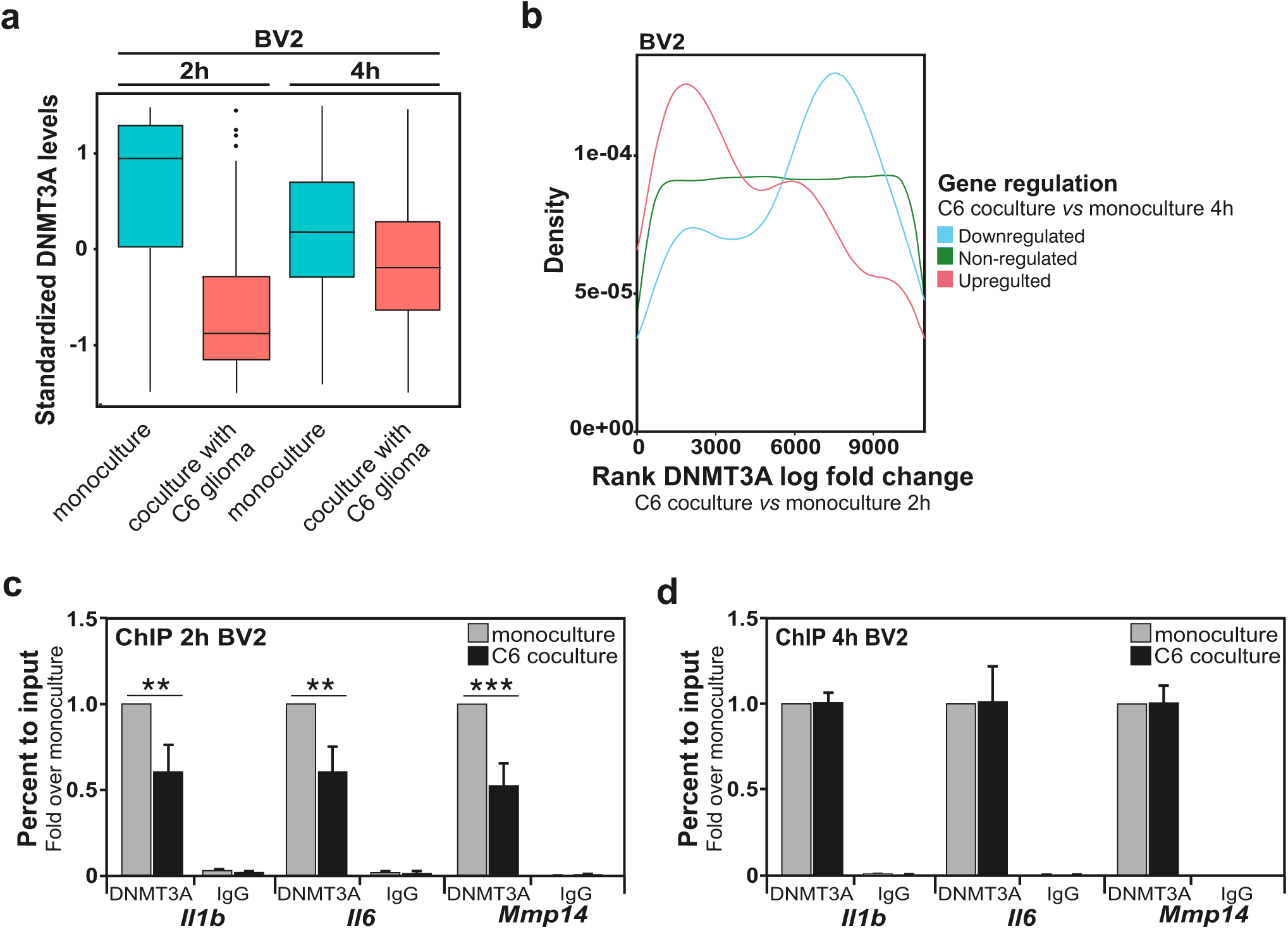
Transient reduction in DNMT3A DNA occupancy is observed in glioma-stimulated microglia. **a**, Box plot representation illustrating at the genome-wide level a general reduction in DNMT3A DNA occupancy in BV2 microglia stimulated for 2 or 4 hours by C6 glioma cells; data obtained from DNMT3A ChIP-seq (n=3). **b**, Analysis of the relationship between DNMT3A DNA occupancy at 2h in C6 glioma-stimulated BV2 microglia (as compared to BV2 alone) and gene expression at 4 hours in C6 glioma-stimulated BV2 microglia (as compared to BV2 alone). The x axis shows the DNMT3A DNA occupancy log fold change ranked from the lowest occupancy (0) to the highest one (∼10000) when BV2 microglia cocultured with C6 glioma compared to BV2 microglia monoculture at 2 hours. Genes were then grouped according to their regulation (3 groups: downregulated, non-regulated, or upregulated) in C6 glioma-stimulated BV2 microglia at 4h (as compared to BV2 alone). The density curves show how those gene groups are distributed in relation to the DNMT3A DNA occupancy. **c**,**d**, ChIP analysis of DNMT3A occupancy on the promoter region of *Il1b*, *Il6* and *Mmp14* genes in BV2 microglia after coculture with C6 glioma cells for 2 hours (**c**) and 4 hours (**d**) compared to their respective BV2 microglia monoculture. Data are presented as percent to input, and fold over monoculture condition set as 1. (n=5 (c), n=3 (d); Data are shown as mean ± sem).

Then, to get further insights into possible role for the displacement of DNMT3A from the chromatin into the regulation of an initial microglial reactive state induced by glioma cells, DNMT3A protein occupancy on the promoter regions of *Il1b, Il6* and *Mmp14* genes (early regulated microglial genes in response to glioma cells stimulation) was analyzed by ChIP in BV2 microglia collected from monoculture or from 2 and 4 hours segregated coculture with C6 glioma cells (Fig.4c,d). A robust decrease in DNMT3A protein occupancy at these 3 gene promoters was observed at the 2 hours’ time-point in microglia collected from coculture with glioma cells, as compared to monoculture conditions (Fig.4c). This difference in DNMT3A protein occupancy between these two conditions was lost at the 4 hours’ time point, indicating a short and transient mechanism for the regulation of these microglial genes by DNMT3A (Fig.4d).

Collectively these experiments show that glioma-induced loss of DNMT3A-mediated DNA methylation at specific gene loci promotes their expression in microglia and contributes to the acquisition of a unique transcriptome associated to a pro-inflammatory and motile microglial cell phenotype.

### *DNMT3A* silencing in microglia promotes a pro-inflammatory/anti-tumoral phenotype

It is interesting to hypothesize that the observed downregulation of microglial DNMT3A and associated transcriptional effects, in response to a glioma stimulus, contributes to the polarization of the microglia cells toward a phenotype that could exert anti-tumoral functions. We therefore decided to assess the role for DNMT3A in the activation of BV2 microglia by knocking down the expression of endogenous *Dnmt3a* using a pool of small interfering RNAs (siRNAs), thereby mimicking the effect glioma cells have on microglial DNMT3A expression. This *Dnmt3a*-targeting siRNAs pool led to robust downregulation of the expression of the *Dnmt3a* mRNA (but did not affect *Dnmt1 and Dnmt3b* mRNAs expression) (Extended Data Fig.4a), as well as the expression of DNMT3A protein in BV2 microglia (Fig.5a). RTqPCR analysis revealed that *Dnmt3a* gene silencing in BV2 microglia was associated with a 2-fold increase in *Il1b and Nos2* mRNA expression levels while *Il6* mRNA expression level was left unaffected (Fig.5b).

**Figure 5.**
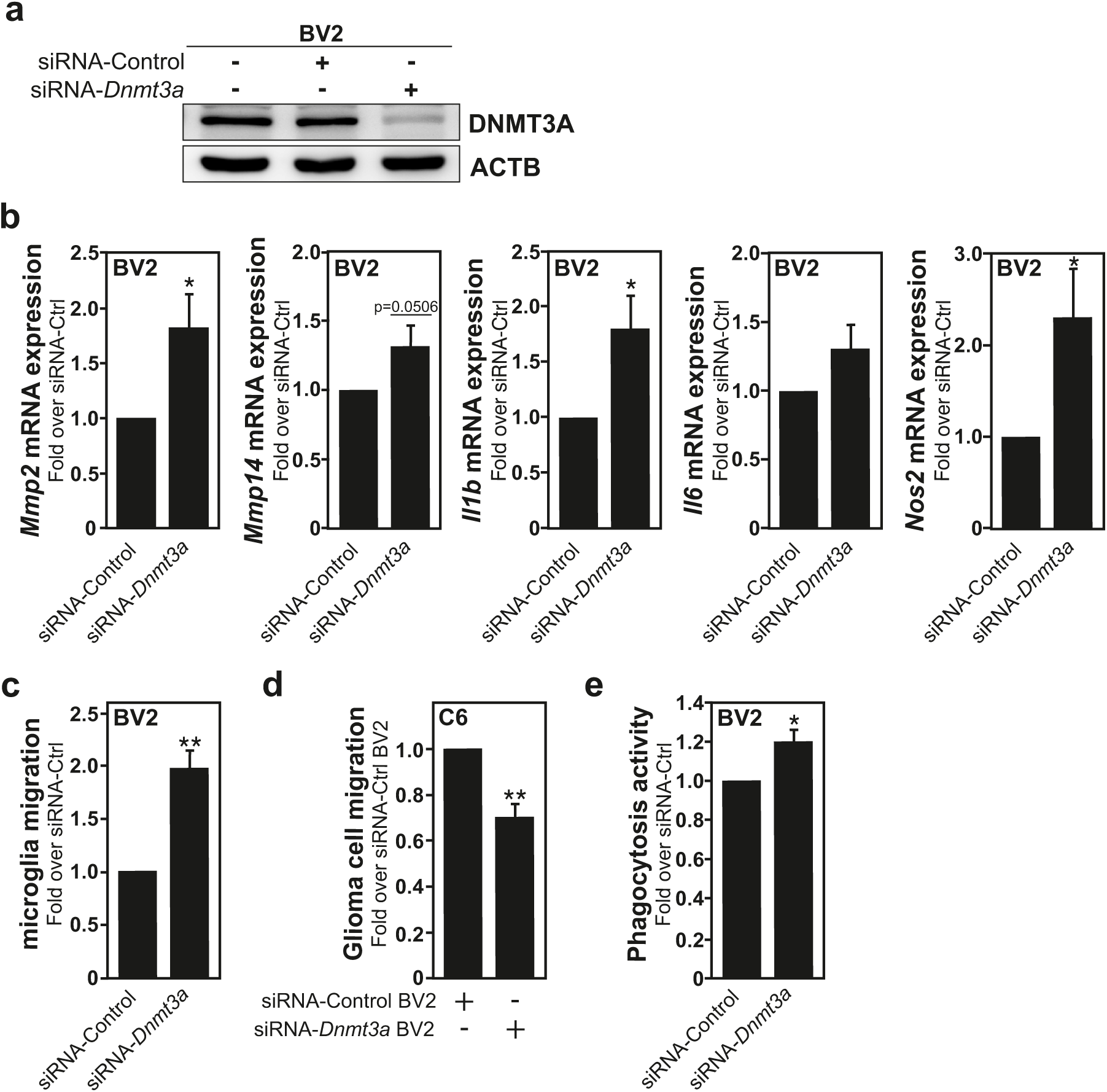
*DNMT3A* silencing in microglia promotes an anti-tumoral phenotype. **a**, Immunoblot analysis of DNMT3A and β-actin (ACTB) expression levels in control BV2 microglia (-, mock transfected cells) or BV2 microglia transfected with a pool of non-targeting siRNAs control (siRNA-Control/Ctrl, negative control) or with a pool of siRNAs targeting *Dnmt3a* expression (siRNA-*Dnmt3a*). **b**, RTqPCR analysis of *Mmp2*, *Mmp14, Il1b, Il6* and *Nos2* mRNA expression levels in BV2 microglia transfected with siRNA-*Dnmt3a* compared to cells transfected with siRNA-Control, set as 1. (n=3-5; Data are presented as mean ± sem). **c**, BV2 microglia transfected with siRNA-*Dnmt3a* exhibit increased cell migration capability, when compared to cells transfected with non-targeting siRNA-Control. **d**, C6 glioma cells migration toward BV2 microglia is reduced when BV2 cells are transfected with siRNA-*Dnmt3a* compared to microglial cells transfected with siRNA-Control. **e**, Phagocytosis activity of BV2 microglia is reduced in cells transfected with siRNA-*Dnmt3a* compared to cells transfected with siRNA-Control. (n=3 (c,d), n=5 (e); Data are presented as mean ± sem).

Then, in a transwell system, we studied the effects microglial *Dnmt3a* gene silencing has on the capability of microglia to migrate, and the ability of these reactive cells to promote the migration of glioma cells. Whereas the inhibition of microglial DNMT3A expression appears to stimulate the migration capability of BV2 microglia cells (Fig.5c), it reduces their aptitude to encourage the migration of C6 glioma cells (Fig.5d). Furthermore, in accordance with their increased migration capability, the gene expression of 2 MMPs, *Mmp14* and *Mmp2* were found to be increased upon repression of *Dnmt3a* expression in the BV2 microglia (Fig.5b). Knockdown of *Dnmt3a* gene expression in BV2 microglia also promoted their phagocytic capacity as revealed by phagocytosis assay with beads (Fig.5e).

Collectively, these observations advocate that the sustained repression of DNMT3A expression in microglia is sufficient to promote an immune and inflammatory phenotype (similar to the one observed in microglia upon short exposure to glioma cells).

### *In vivo,* DNMT3A-deficient microglia reduce glioma tumor growth

Based on our finding that the regulation of DNMT3A is involved in the control of a gene expression profile in microglia associated with inflammatory and immune reactive responses, which can potentially exert anti-tumoral functions, it is attractive to speculate that repressing its expression in microglia could provide us with cells which exert beneficial, anti-tumoral effects in the context of glioma tumors.

Viral delivery of small hairpin RNA (shRNA) targeting *Dnmt3a* was used for the establishment of BV2 microglia-derivatives with stable knockdown of DNMT3A expression (Extended Data Fig. 5a,b). A *Dnmt3a* shRNA expressing microglial clone (*i.e.,* BV2 shRNA *Dnmt3a*#2) that exhibited a 90% reduction in DNMT3A protein expression, as compared to a microglial clone infected with the lentiviral empty vector control shRNA (shRNA Control), was selected for further experimentation. To examine the physiological relevance of our findings, we performed *in vivo* experiments and injected GFP-expressing GL261 glioblastoma cells or co-injected GL261 cells together with BV2 microglia expressing a *Dnmt3a* shRNA into the brain of young C57BL6/J mice. Tumors established from GL261 cells recapitulate numerous GB characteristics.^44^ Importantly, this syngeneic transplant tumor model in immunocompetent mice has been shown to exhibit limited infiltration by peripheral monocytes or macrophages at the time points used.^11^ Immunohistochemical analysis of brain tissue 2 weeks post-transplantation revealed a trend, yet not significant, for a reduction of GFP-GL261 glioma tumor sizes in mice co-injected with the BV2 microglia expressing *Dnmt3a* shRNA as compared to those injected with the GFP-GL261 alone (Extended Data Fig.5c,d), suggesting that repression of DNMT3A expression in microglia could induce an effective anti-tumoral phenotype in these myeloid cells.

Encouraged by these promising results, and to gain direct evidence that reducing microglial *Dnmt3a* expression *in vivo* could lead to reduction in GB tumor growth, we made use of the antisense oligonucleotides (ASOs) developed by Ionis^®^ Pharmaceuticals to target *Dnmt3a* in a highly specific manner. RNA antisense therapies, including for CNS disorders (*e.g*., Ulefnersen, Tofersen, Zilganersen, ION859/BIIB094), have shown promising preclinical settings as well as in clinical trials (ClinicalTrials.gov).^45–47^ Two different mouse-specific *Dnmt3a*-targeting ASOs (referred to as ASO-1 and ASO-2) were developed and tested *in vivo*. Thereafter, the efficiency of these ASOs was tested at 2 and 8 weeks after injection in the right ventricle of C57BL6/J mice brains and showed strong reduction in global *Dnmt3a* expression into the cortex and spinal cord of the injected mice (Extended Data Fig.6a,b). Both mouse-specific *Dnmt3a*-targeting ASOs were found to be safe based on delayed functional observational battery (FOB) and RTqPCR analysis of neuroinflammation markers. Further analyses using RNAscope and immunofluorescence confocal imaging confirmed the reduction of *Dnmt3a* mRNA and DNMT3A protein levels 2 weeks post-injection of either ASO-1 or ASO-2, as compared to a control non-targeting ASO (ASO-Ct) (Fig.6a,b).

**Figure 6.**
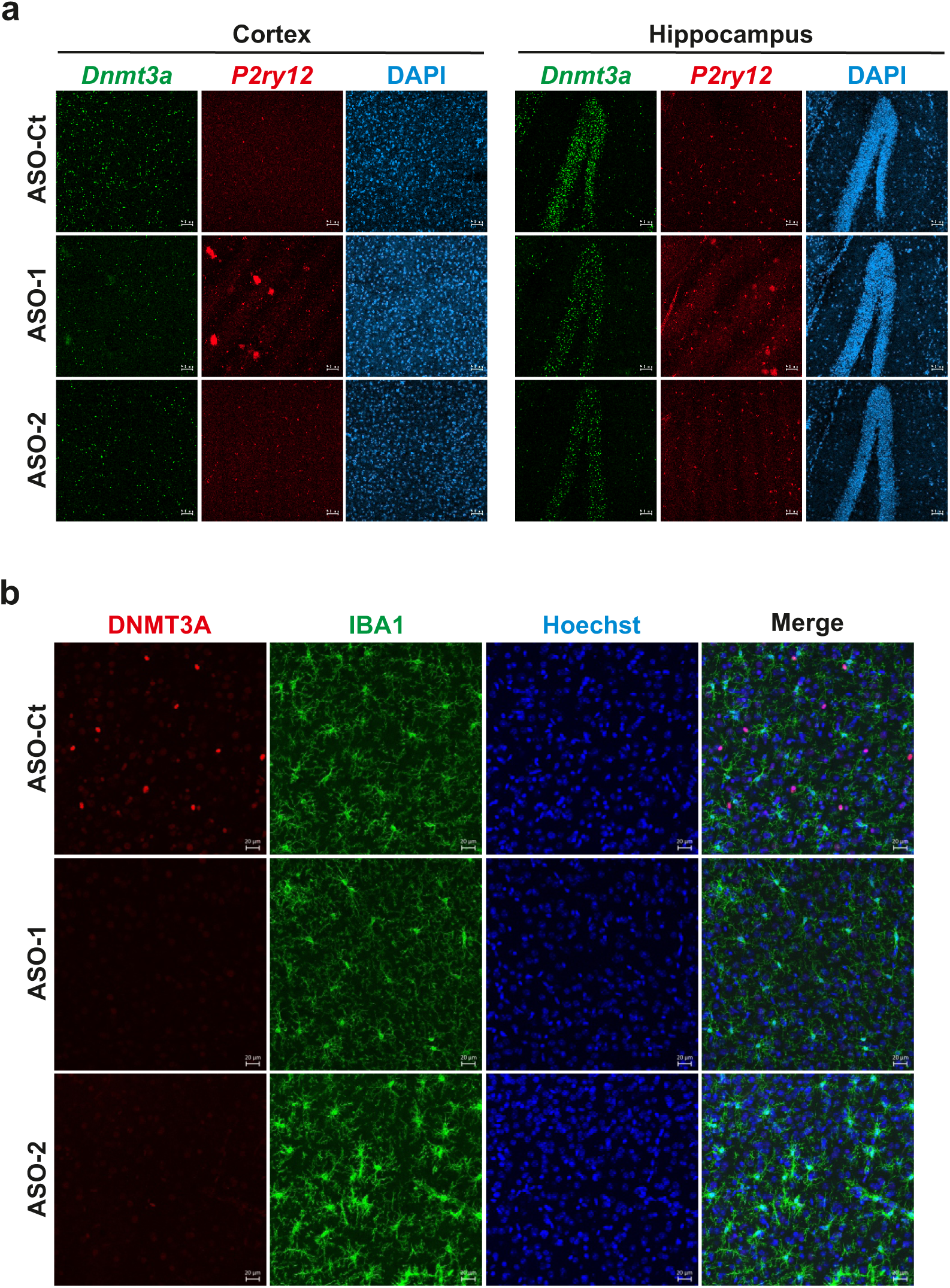
ASOs targeting *Dnmt3a* efficiently reduce *Dnmt3a* mRNA and DNMT3A protein expression levels *in vivo*. **a**, Confocal analysis of *Dnmt3a* (green) and *P2ry12* (red, used as a microglial marker) mRNA expressions by RNAscope in the cortex (left panel) and hippocampus (right panel) brains regions of mice 2 weeks post injection of non-targeting ASO (ASO-Ct) or ASOs targeting *Dnmt3a* (ASO-1 and ASO-2). DAPI (blue) was used as nuclear counterstain. (Scale bar, 50µm) **b**, With similar conditions as described in panel a, confocal immunofluorescence analysis of DNMT3A and IBA1 (used as a microglial marker) protein expressions (red) in mice brains. Hoechst (blue) was used as nuclear counterstain. (Scale bar, 20µm).

Analysis of microglia in the brains of mice injected with ASO targeting *Dnmt3a* highlighted that the reduction of *Dnmt3a*/DNMT3A expression induces a microglial reactive state. Indeed, microglia in brains injected with the ASO-2 displayed an amoeboid shape, characteristic of activated microglial cells, associated with a significant increase of ionized calcium binding adaptor molecule 1 (IBA1, also known as Allograft inflammatory factor 1, AIF1) immunofluorescence intensity (Fig.6b and Fig.7a,b). Additional brain immunofluorescence confocal imaging revealed an increase in staining intensity for Galectin-3 (LGALS3), whose expression is associated with a microglial inflammatory response^48^, 2 weeks after the injection of ASO-2 compared to ASO-Ct; further validating the presence of an immune activated microglial phenotype after *Dnmt3a* knockdown (Extended Data Fig.6c).

**Figure 7.**
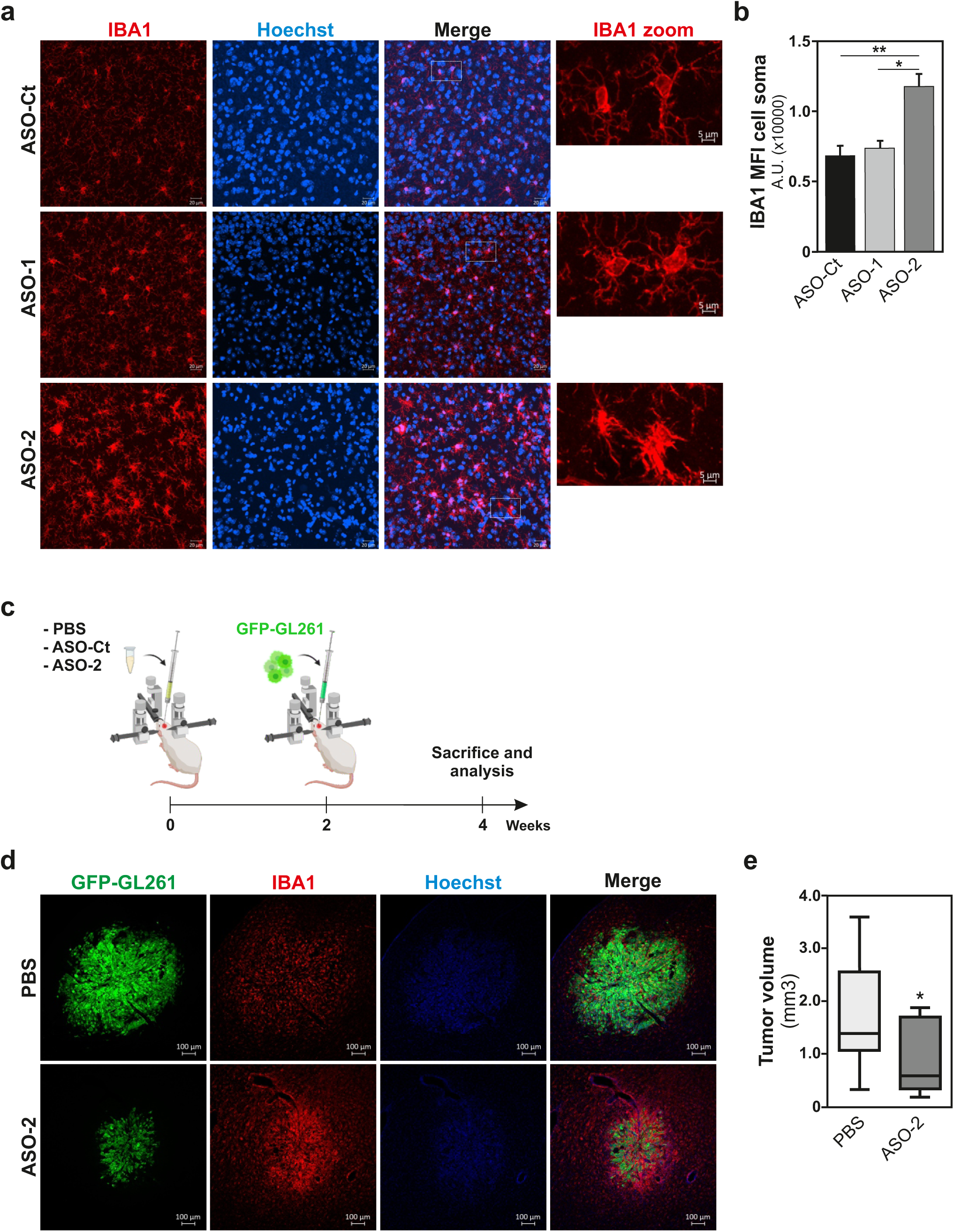
*Dnmt3a* knock-down in microglia induces microglia activation and decreases glioma tumor growth *in vivo*. **a**, Confocal immunofluorescence analysis of microglial IBA1 expression (red) and Hoechst-nuclear couterstaining (blue) in brain tissues of mice injected with ASO-Ct or ASO targeting DNMT3A (ASO-1 and ASO-2). The white rectangles indicate the regions magnified and presented on the right-hand side of each condition, that illustrate microglia harboring different shape depending on the treatment (referred to as IBA1 zoom). (Scale bars, 20µm, 5µm in zoom). **b**, Quantification using IMARIS software of the IBA1 intensity (mean per cell) in microglia from mice treated with a non-targeting control ASO (ASO-Ct) or ASOs targeting *Dnmt3a* expression (ASO-1 and ASO-2). **c**, Scheme representing the workflow of the *in vivo* experiments performed with the GFP-GL261 GB mouse model presented in panels d and e. **d**, Confocal immunofluorescence imaging of tumors formed 2 weeks post injection of GFP-GL261 glioma cells (green) in mice previously treated with PBS, or ASO-2 (as illustrated in c) together with an immunostaining for the microglial marker IBA1 (red) and Hoechst used as nuclear counterstain (blue). **e**, Quantification of 2 weeks old GL261 tumor volume in mice injected with PBS or ASO-2, 2 weeks prior injection of GFP-GL261 cancer cells. (n=9/10 mice per group; Data are presented as box plot)(Scale bars 100µm).

Finally, we investigated the potential anti-tumoral properties of our ASO targeting *Dnmt3a* in an *in vivo* GB tumor mouse model. Based on the above observations that *in vivo* the ASO-2, as compared to ASO-1, i) had a better acute tolerability, and ii) was more effective in promoting a microglial reactive state, this *Dnmt3a*-targeting ASO was selected for further *in vivo* investigations. Two weeks prior to the transplantation of GFP-GL261 GB cells into the brain of C57BL6/J mice, those mice were injected with ASO-2 targeting *Dnmt3a* expression, or with PBS, *i.e*., vehicle for ASO injection, used as control (workflow in Fig.7 c). Two weeks post-injection of the GFP-GL261 GB cells, mice were sacrificed, and the presence of microglia as well as tumor size were investigated by immunofluorescence confocal imaging (Fig.7d). IBA1 used to detect the microglial cell population was observed to be more prominently expressed in the brains of ASO-2 injected mice compared to PBS-injected ones (Fig.7d). In addition, detection of the GFP-GL261 GB cells after ASO treatment revealed a significant reduction in tumor growth in mice injected with the ASO-2 compared to PBS-injected animals (Fig.7d,e). Importantly, the injection with the ASO-Ct did not yield the effects observed with ASO-2 on the GFP-GL261 GB tumor growth (Extended Data figure 6d,e).

In summary, specific ablation of microglial DNMT3A negatively affects their tumor-supporting function in favor of anti-tumoral properties, and an RNA-based antisense strategy that counteracts selectively *Dnmt3a* expression led to both microglia activation and reduced glioma expansion *in vivo*.

## Discussion

A widespread view in the literature is that glioma cells recruit microglia, resident immune cells within their brain microenvironment, and convert them into tumor-supporting cells. There are numerous glioma cells-derived factors that can promote the recruitment of microglia toward a tumor site, including chemokines, cytokines, ligands of complement receptors, neurotransmitters and ATP.^2, 15, 32, 49^ During that process, the tumor cells are reported to shut down their immune properties and stimulate the microglia to exert instead tumor cell trophic effects, thereby contributing to a pro-invasive and immunosuppressive tumor environment. In fact, microglia release many factors, including extracellular matrix metalloproteinases and cytokines, which directly or indirectly influence tumor invasiveness and growth.^13, 16, 50, 51^ At the transcriptomic level, glioma cells are reported to hijack microglial gene expression to promote tumor growth and expansion.^3, 4, 18^ In murine GB models, advanced tumor (*e.g*., 4 weeks post intracranial injection of GL261 GB cells in C57BL/6 mice brains, syngeneic transplant in an immunocompetent environment) are characterized by the downregulation of the expression of microglial genes involved in the sensing and dismissal of tumor cells. At the same time, the microglial expression of genes involved in facilitating tumor spread and expansion are upregulated.^3, 4, 18^ Likewise, the analysis of transcriptomic data sets obtained from human GB biopsies, reveal a proliferative and anti-inflammatory phenotype for the tumor-associated microglia that preferentially reside at the leading edge of tumor infiltration.^7, 8, 52, 53^

Our findings challenge this view, as it appears that glioma cancer cells initially promote a microglial phenotype which is associated with a gene expression profile associated to both pro-inflammatory and immune responses. However, this immune responsive phenotype is temporary, and the microglia, in response to cancer cells stimulation, turn into persistent tumor-supportive cells. Supporting the existence of a transient glioma cancer cells-induced microglial immune reactive state, in segregated cocultures performed with human CHME5 or murine BV2 microglial cells together with murine C6 glioma cells, transitory modifications of microglial morphology and metabolism were observed and found to be linked to a concomitant transitory increase of phagocytic capability; properties that got lost after longer exposure to glioma stimulation.^54^ Further evidence comes from *in vivo* observations made with the GL261 GB mouse model, at times where microglia recruitment and tumor growth are observed (*i.e*., 1 and 2 weeks post brain implantation of the GL261 GB cells).^11, 12^ Indeed, at those earlier stages of GB development, microglia outside the tumor mass were shown to express the active caspase-3 that has been link to a microglial proinflammatory phenotype, whereas microglia found within the core of the tumor showed an inhibition of basal caspase-3 activity associated with the polarization of microglia toward a tumor-supportive phenotype.^19, 33^ Hence, delineating the molecular mechanisms that control the microglial activation via a transient, yet potentially anti-tumoral phenotype, in response to cues from glioma cells is of considerable interest.

The cellular reprogramming of microglia toward a particular phenotype with unique functions requires the acquisition of a new gene expression signature, in the context of unchanging genomic sequence. Epigenetic changes, including histone modifications, microRNA expression, as well as DNA methylation, are important modulators of gene expression, and have been involved in cell phenotype regulation and reprogramming, and are therefore part of the mechanisms regulating cellular plasticity. In contrast to the extensive investigations made on the effects of histone modifying enzymes and associated histone posttranslational changes, or of microRNA on microglial gene expression, the regulation of gene expression by DNA methylation remains poorly explored in those cells (for review see ^20, 55^).

DNA methylation catalyzed by DNMTs is a relatively stable, yet reversible, epigenetic modification that plays important roles in the establishment and stabilization of cellular phenotypes by maintaining gene expression profiles.^21^ In fact, DNA methylation can be conserved during cell division, and is mostly associated with sustained transcriptional repression. This heritable chromatin mark has been involved in various cell reprogramming processes.^22^ Hence, DNA methylation mediated by DNMTs is often viewed as mediator of long-term epigenetic effects. However, we uncovered that glioma-induced microglia activation is associated with the transient displacement of microglial DNMT3A from chromatin sites, reduction in DNA methylation, and the induction of the expression of genes that confer to these cells a temporary transcriptome profile reflecting an immune reactive phenotype. Worth a notice, this distinct microglial DNMT3A-mediated signaling pathway appears to be induced in response to exposure to glioma cells, but not upon stimulation with IL4, a treatment frequently used to mimic some of the effect of glioma cells on microglial cells, or upon stimulation with LPS, used to initiate a pro-inflammatory response in microglia.

The involvement of a prompt but transient reduction of DNA methylation in the reprogramming of microglia toward a temporary state of activation, may be seen as unusual epigenetic stress response, however similar mechanism has been reported earlier on for other cell types undergoing differentiation.^56–59^ In plants, active DNA demethylation regulates many developmental processes as well as their response to various environmental stress.^60, 61^ In mammalian cells, active DNA demethylation also contribute to cell plasticity. For example, a transient DNA demethylation driven by a loss of DNA methyltransferase proteins is a molecular characteristic of the mouse embryonic stem cells subpopulation which cycle through the so-called MERVL/Zscan4 cell state.^59^ In addition, active DNA demethylation appears to be a common feature of the response of human innate immune cells, *i*.*e.*, dendritic cells, T cells, natural killer cells, neutrophils, and macrophages, facing an infectious agent.^62–66^ Our discovery that in response to glioma stimulation, a transient DNA demethylation driven by reduced DNMT3A chromatin occupancy, is associated with the induction of gene expression profile reflecting both pro-inflammatory and immune responses, and thus potentially anti-tumoral functions in microglial cells, argue for the targeting of this microglial DNA demethylation-mediated signaling pathways for therapeutic purpose to treat gliomas.

DNA demethylating agents, de facto DNMTs inhibitors, such as Azacytidine and Decitabine (5-aza-2′-deoxycytidine), in combination with temozolomide, radiation or even oncolytic virus, have been shown to hold beneficial effects in *in vitro* and *in vivo* glioma models.^67–71^ However, despite these pre-clinical tests and a few clinical trials (see clinicaltrials.gov) DNA demethylating agents have failed to establish themselves as part of the treatment protocols for high-grade gliomas. Therefore, more targeted strategies in terms of both the molecular and the cellular targets should be considered. Our findings that glioma cells induce a significant decrease in DNMT3A expression in microglia that in turn hold potential anti-tumoral properties, urges to investigate whether microglial DNMT3A can be targeted selectively to treat glioma.

RNA-based antisense technology is reported to be an efficient method to decrease the expression of selective DNMT protein^72, 73^, and this approach has even attracted interest in the gliomas’ research field.^74^ In addition, microglia have been considered to be interesting cellular target to treat glioma, however recent efforts have been focused on repressing their tumor-supporting functions, or even achieving their depletion (for review see ^75^). Instead, we propose to capitalize on the in-built anti-tumoral properties of microglia, which we now discovered can be stimulated by the cancer cells themselves.

Our investigations reveal that i) the expression of DNMT3A can be selectively and robustly reduced in microglia using RNA-based antisense approaches, transiently using small interfering RNA and even in a stable manner using small hairpin RNA, ii) that reduced DNMT3A expression is associated with a microglial phenotype holding anti-tumoral properties, and finally iii) that targeting microglial *Dnmt3a* using ASOs impacted negatively on the growth of GB tumors in an animal model of the disease. Thus, taking advantage of a glioma-induced microglial signaling pathway that controls the acquisition of a transient microglial phenotype with inflammatory and immune responses characteristic, we can treat the neoplastic cells. Furthermore, the modulation of microglia DNMT3A expression could provide an option to override the immune inhibitory and tumor-supportive functions exerted by these cells in the context of the glioma tumor microenvironment and should therefore be further considered as therapeutic strategy.

## Material and methods

### Cell culture

BV-2 (referred as BV2) mouse microglia cell line (RRID:CVCL_0182; gift of G. Brown, University of Cambridge)^19, 33^, HMC3 human microglia cell line (RRID: CVCL_II76; also known as CHME3, obtained from originator M. Tardieu, Paris-Sub University)^76^, GL261 mouse glioblastoma cell line expressing the enhanced green fluorescent protein (GFP) (referred as GFP-GL261) (RRID:CVCL_Y003; gift of R. Glass, Max Delbrück Center)^13^ and C6 rat glioblastoma cell line (RRID:CVCL_0194; gift of O. Hermanson, Karolinska Institutet) were cultivated in DMEM + glutamax medium (Gibco) supplemented with 10% FBS (fetal bovine serum) and 1% P/S (Penicillin/Streptomycin). U-87MG ATCC (referred as U87) human glioblastoma cell line (RRID: CVCL_0022; purchased from ATCC, HTB-14™) was cultivated in MEM medium supplemented with 10% FBS and 1% P/S. All cell lines were grown in an incubator at 37°C and 5% CO2 and regularly tested with Venor GeM mycoplasma detection kit (Minerva Biolabs).

For segregated coculture experiments, microglial cells were first seeded in 5% FBS medium on coverslips in 12 well plates while glioblastoma cells were seeded in 94 mm diameter Petri dishes. Thereafter, 24 hours after seeding, the cocultures were initiated by placing the coverslips with microglial cells into cell strainers in the petri dishes containing the glioblastoma cells. Microglial cells were then harvested at given time-points.

### Gene silencing by transfection of small interfering RNAs pools

For transient gene expression silencing by siRNAs, non-targeting control, *Dnmt1*, *Dnmt3a* and *Dnmt3b* ON-TARGET plus SMARTpools siRNAs, whose sequences can be found in the Supplementary Table 1, were obtained from Dharmacon. Transfection of BV2 cells was carried out with Lipofectamine 3000 (Invitrogen).

### Dnmt3a gene silencing by short hairpin RNA lentiviral infection

For stable gene expression silencing of *Dnmt3a* by shRNA, MISSION^®^ shRNA for *Dnmt3a* in pLKO-PURO vector in lentiviral particles, and an empty pLKO-PURO vector used as control, were purchased from Sigma Aldrich. Two clones (Clone IDs: TRCN0000039034 with target sequence: CCAGATGTTCTTTGCCAATAA, and TRCN0000039035 with target sequence: GCAGACCAACATCGAATCCAT) were tested for lentiviral infection at different Multiplicity of Infection (MOI 1, MOI 2 and MOI 5) of the BV2 cells overnight in the presence of polybrene (Sigma-Aldrich). The MOI of 2 was selected as the best ratio of infection. Two days post-infection, cells that incorporated the shRNA were selected using fresh medium containing 5µg/ml of Puromycin (Sigma-Aldrich) and shRNA knockdown efficiency was monitored by immunoblot analysis (as described below) after the first 3 cell passages and at regular intervals when running experimental work.

### Immunoblotting

Total protein extracts were made directly in Laemmli buffer by scraping of the cells. For immunoblot analysis, protein extracts were resolved on 8% SDS–polyacrylamide gel electrophoresis and then blotted onto 0.45 µm pore-size nitrocellulose membranes. Membranes were blocked with 0.1% Tween/5% milk in PBS and incubated with indicated primary antibodies, overnight at 4°C, followed by incubation with the appropriate horseradish peroxidase secondary antibody (Pierce, 1:10,000) for 1 hour at room temperature 20-25°C. Immunoblot with anti-β-actin antibodies was used for standardization of protein loading. Details about antibodies used in this study can be found in Supplementary Table 2. Bands were visualized by enhanced chemiluminescence (ECL-Plus, Pierce) following the manufacturer’s protocol. Alternatively, the membranes were incubated with the appropriate secondary antibody, RDye^®^ 680RD Goat anti-Rabbit or RDye^®^ 800CW Goat anti-Mouse IgG (1:5000; LI-COR Bioscience, Lincoln, NE, USA) for 1h at room temperature and visualized using an Odyssey CLx infrared imaging system (LI-COR Bioscience). All targeted proteins of interest were normalized to the selected housekeeping gene, intensity of the bands was verified within the same linear range and quantification was performed using the ImageJ software.

### DNA extraction

DNA extraction was performed using the QIAamp DNA Mini Kit (Qiagen). DNA concentration was quantified using NanoDrop^®^ spectrophotometer (Thermo Fisher Scientific).

### DNA methylation analysis

Whole genome DNA methylation analysis was performed using the Illumina^®^ Infinium HumanMethylation450 BeadChip array. Experiments were performed according to protocol at the Bioinformatics and Expression Analysis (BEA) Core facility at Novum, Karolinska Institutet.

### Dot Blot

500 µg of DNA in 0.1M NaOH (or positive 5mC DNA used as control, Diagenode) were denaturated 5 min at 99°C. The samples were then neutralized with 0.1 volume of 6.6M ammonium acetate and spotted on a Hybond-N+ membrane (Amersham). The membrane was air dried before cross-linking 2 hours at 80°C. The membrane was then blocked in 10% milk, 1% BSA diluted in PBS-tween (0.1%) for 1 hour at room temperature before the overnight incubation at 4°C with the primary antibody directed against 5-methylcytosine (Diagenode) diluted 1/250 in blocking solution. After 3 washes in PBS-tween the appropriate horseradish peroxidase secondary antibody (Pierce, 1:10,000) was incubated for 1 hour at room temperature. Spots were visualized by enhanced chemiluminescence (ECL-Plus, Pierce) following the manufacturer’s protocol. Densitometry was done using the ImageJ software.

### RNA isolation, cDNA synthesis, and qPCR

Total RNA was extracted using the RNeasy Mini Kit (Qiagen). RNA concentrations were quantified using NanoDrop® spectrophotometer (Thermo Fisher Scientific). cDNA was synthesized from 1 µg RNA using Oligo dT, dNTPs, and Superscript III Reverse Tanscriptase (Invitrogen). qPCR was run on StepOne plus (Applied Biosystems) using the SYBR™ Green master mix (Life Technologies) and primers listed in Supplementary table 3. *Actb* gene expression in each sample was used for normalization. Results were calculated using the Δ*C_t_* method and represented as a fold over control (microglia cells from monoculture condition).

### RNA sequencing

Total RNA was subjected to quality control with Agilent Tapestation according to the manufacturer’s instructions. 200 ng of Total RNA was subjected to Illumina sequencing and libraries were prepared with the Illumina TruSeq Stranded mRNA kit which includes cDNA synthesis, ligation of adapters and amplification of indexed libraries. The yield and quality of the amplified libraries was analysed using Qubit by Thermo Fisher and the Agilent Tapestation. The indexed cDNA libraries were normalized and combined, and the pools were sequenced on the Illumina Hiseq 2000 generating 50 bp single-end reads. Basecalling and demultiplexing was performed using BCL2 software with default settings generating Fastq files for further downstream mapping and analysis.

### Transcriptome data processing

The top 100 most significant Gene Ontology (GO) pathways found in the upregulated genes between co and monocultured cells using the weight01 algorithm were taken and the semantic similarity was calculated pairwise between the GOs using the relevance method. The GOs were then clustered using hierarchical correlation clustering. The color shows the degree of similarity and the dendrograms the clusters formed based on their similarity.

### Computational analysis

Differentially expressed genes with significant (< 0.05) FDR were used for all further analysis. The genes shown in the heatmap are genes that were found significantly regulated in any of the comparisons. Each gene (row) is standardized (z) to mean = 0 and sd = 1 and then clustered by hierarchical clustering. For the ‘positive versus negative’ comparison, a volcano plot was created in GraphPad Prism, which displayed significance and FC for the dataset together with gene symbols for the most highly regulated genes. The analysis of Gene Ontology (GO) terms was performed using the bioinformatics database Metascape (https://metascape.org/, 77). Gene lists of interest were imported and the resulting enrichment map file then adjusted for visual preferences using the Cytoscape software (v3.7.1, www.cytoscape.org; 78). The Enrichment Map (www.baderlab.org/Software/EnrichmentMap) plug in was used as a visualization tool to produce a network graph of over-represented GO terms. Enrichment was based on an FDR of maximum 0.05. Nodes represent enriched GO terms and node size correlates with the number of genes included in a specific GO term. Edges indicate the degree in overlap between nodes. In addition to the enrichment maps for all significant differentially expressed genes, enrichment analysis was performed on a subset of those genes that overlap with immunologic signature genes.

A heat map for validated genes found to be differentially expressed (up- or downregulated by at least twofold) between the two microglial populations of interest was generated using GraphPad Prism.

### Chromatin immunoprecipitation (ChIP) and ChIP sequencing

DNMT3A ChIP experiments were done using the HighCell# ChIP kit from Diagenode (kch-mahigh-G48) according to the manufactureŕs instructions. Briefly, after cell cross-linking in 1% formaldehyde and cell lysis, chromatin shearing was done with a Bioruptor® Pico sonicator (Diagenode). Then, each chromatin immunoprecipitation was done using 6.3μg of antibody. Purified DNA and 1% input were then analyzed by qPCR (primers listed in supplementary Table 2). Data interpretation from qPCR was done by calculation of the percentage to input and then normalized to the control condition.

DNMT3A ChIP-seq experiments were done using the iDeal ChIP-seq kit from Diagenode according to the manufactureŕs instructions. Purified DNA was sent for library preparation and sequencing at the Bioinformatics and Expression Analysis core facility (BEA, Novum, Karolinska Institute).

ChIP DNA was subjected to quality control with Agilent Tapestation according to the manufacturer’s instructions. To construct libraries suitable for Illumina sequencing the NEB Ultra DNA kit was used. 10 ng of chipped DNA was used as input. The protocol includes ligation of adapters and amplification of indexed libraries and purification with AMpure magnetic beads. The yield and quality of the amplified libraries were analyzed using Qubit by Thermo Fisher and the Agilent Tapestation. The indexed DNA libraries were normalized and combined, and the pools were sequenced on the Illumina Hiseq 2000 for a 50-cycle sequencing run generating 50 bp single-end reads. Basecalling and demultiplexing was performed using CASAVA software with default settings generating Fastq files for further downstream mapping and analysis.

### Transwell migration assay

Eight μm-pore width transparent PET membrane inserts (Transwell, Corning) were used to measure cell migration capability using a transwell system. For C6 glioma cells migration assay, C6 cells were seeded on top of the insert and BV2 microglia were seeded in the lower compartment. For BV2 microglia migration assay, BV2 cells were seeded in the insert in 5% FBS medium and 10% FBS medium was placed in the bottom as attractant. Once the experiment was finalized (after 24 hour migration for C6 cells and 4 hour migration for BV2 cells), the membranes from the inserts were washed with PBS and carefully cut out with a blade. The membranes were mounted with ProLong Gold antifade reagent with DAPI (Life technologies) and the nuclei of the migrated cells were counted under fluorescent microscopy.

### Syngeneic transplant glioma mouse model

Experiments were performed in accordance with the Guidelines of the European Union Council, following Swedish regulations for the use of laboratory animals and approved by the Regional Animal Research Ethical Board, Stockholm, Sweden (Ethical permits N248/13, 17283/2018, and 13676/2020). C57/BL6/J mice (Charles River) were housed in a 12 hour light/12hour dark cycle with access to food and water ad libitum.

#### Intrastriatal GL261 and BV2 injections

Postnatal day 16–17 male pups were anesthetized with isoflurane (5% for induction and 1.5% for maintenance). An incision was made on the scalp and the skin flaps were retracted to expose the skull. Animals received an intrastriatal injection of either Gl261 alone (35,000 cells) or a mix of GL261 and BV2 shDNMT3A (35000 + 15000 cells respectively) resuspended in 1 μl of the culture medium in the left hemisphere using the following coordinates relative to bregma anterior/posterior: +0.7 mm, lateral: ± 2.5 mm, ventral: −3 mm, using a 5 μl ILS microsyringe. The injection was performed over 1 min and the syringe remained in the injection site for 5 min to reduce backflow and was slowly retracted over 1 min thereafter. The skin was sutured, and animals were allowed to recover before they were returned to their dams. Animals were sacrificed 2 weeks after transplantation. The experiment was performed on 4 animals per condition and repeated twice (n=8 animals per condition final).

#### Antisense oligonucleotide and cortical GL261 injections

Six-week-old mice were anesthetized with isoflurane (5 % for induction and 2 % for maintenance) using a mouse mask coupled to a stereotaxic apparatus. Mice were also injected subcutaneously (s.c.) with 5 mg/kg rimadyl (Carprofen) and 0.1 mg/kg buprenorphine (Temgesic) 15 min prior to surgery for systemic anesthesia and analgesia. The fur on top of the head was removed using a shaver, and the skin was disinfected with ethanol. A 2 mm incision was then made on the scalp following lidocaine administration (4mg/kg). ASOs (ASO-2, or ASO-Ct) or PBS were injected intracerebroventricularly (icv) using the following stereotaxic coordinates: 0.3 mm anterior to bregma (AP), 1.0 mm lateral to bregma (ML), and 3 mm deep (measured from when the orifice of the needle has passed the skull) (DV) using a Hamilton 10 μl syringe (#80030 1701 RN) and a (22s/51/2)S needle (#7758-03 RN) without drilling the skull. The volume (10 μl) and speed (0.1 μl/sec) of the injection were controlled by an automatic microinjection pump. The syringe remained in the injection site for 3 min to reduce backflow and was slowly retracted thereafter. After the surgery, 2-3 stitches (Ethilon monofilament 5.0) were applied to suture the wound. Eye gel drops (Viscotears 2 mg/ml, # 541760) were used to hydrate the eyes throughout the surgery. Mice (n=12/group) were allowed to recover in a separate cage on a heating pad (35.5°C) and monitored until they were fully awake and active. They were also given analgesics for post-operative pain relief (Carprofen 5mg/kg) 24 hours later.

Two weeks after the first injections, mice were again anesthetized with isoflurane and administered rimadyl and buprenorphine s.c. as before. A small craniotomy of 2 mm was performed on the skull using a drill at 1 mm anterior to bregma, 1.5 mm lateral from bregma, and 2.5 mm deep (measured from when the orifice of the needle has passed the hole). A suspension of GL261 tumor cells in PBS (50,000 GL261 tumor cells in 3 µl) was injected in all animals using a 10 µl Hamilton syringe (#80030 1701 RN) and a (26/30/4)S needle (#7804-03 RN) at a speed of 1.5 µl/min. The syringe remained in the injection site for 3 min to reduce backflow and was slowly retracted thereafter. The skin wound was sutured, and the animals were allowed to recover in a separate cage on a heating pad and monitored until they were fully awake and active. They were given analgesics for post-operative pain relief (Carprofen 5mg/kg, s.c., 24 hours later). Animals were sacrificed 2 weeks after cell transplantation.

#### Brain tissue extraction, fixation, and sectioning

Animals were deeply anesthetized with an overdose of sodium pentobarbital and transcardially perfused with 0.9 % sodium chloride, followed by fixation with 4 % paraformaldehyde in 0.1 M phosphate buffer (pH 7.4). Brains were then transferred to 30 % sucrose in 0.1 M phosphate buffer and left until they sank. 25-μm-thick horizontal free-floating sections were prepared using a sliding microtome (Leica SM2010R) and stored in cryoprotection solution at 4°C (25 % glycerol, 25 % ethylene glycol in 0.1 M phosphate buffer) for further histological analysis.

### RNAscope in situ hybridization

RNA *in situ* hybridization was performed for *Dnmt3a* and *P2ry12* mRNAs. After sacrificing the mice, the brains were immediately frozen on dry ice for 5 min and then embedded in cryo-embedding medium (OCT compound, Sakura Finetek). After equilibrating at -20°C for 2 hour in a cryostat (CryoStar NX70, ThermoScientific), the brains were sectioned into 10 μm slices and mounted onto SuperFrost Plus adhesion slide (Epredia). The slices were dried at -20°C for 1 hour and then stored in zip-lock bags at -80°C until needed. The RNAscope Multiplex Fluorescent Reagent Kit (Advanced Cell Diagnostics) for *in situ* hybridization assay was used to detect the mRNAs of interest. The slides were fixed by immersion into pre-chilled 4% PFA for 1 hour at 4°C. After rinsing the slides twice with 1X PBS to remove the fixative, the sections were dehydrated in 50%, 70%, 100%, and 100% ethanol each for 5 min at room temperature. RNAscope hydrogen peroxide was added to each slice for 10 min at room temperature and then washed off with distilled water followed by protease IV for 10 min at room temperature and then washed with PBS. *Dnmt3a* (GenBank accession number NM_153743.4; target nt region, 3757–4994) and *P2ry12* (GenBank accession number NM_027571.3; target nt region, 739 - 1854) probes were pre-warmed for 10 min at 40°C. The probes targeting RNAs contain different 20 ZZ oligonucleotide probes: Dnmt3a-C1 probe, P2ry12-C3 probe. To hybridize with target RNAs, the probes were mixed and added on the slides for 2 hour at 40°C in an HybEZ oven. The signals were then amplified by AMP1, AMP 2, and AMP 3 hybridization for 30 min at 40°C followed by HRP-C1 for 15 min at 40°C and incubation with Opal^TM^ 570 for 30 min at 40°C for *Dnmt3a* detection. The slides were then blocked with HRP blocker before adding HRP-C3 and Opal^TM^ 690 for detecting *P2ry12*. DAPI was used for 30 s at room temperature before mounting the slides with Prolong Gold Antifade Mountant. Pictures were taken using a Zeiss LSM900-Airy confocal microscope and analysed with OMERO.insight and ImageJ software.

### Immunohistochemistry

The mice brain free floating sections were washed in PBS and followed by antigen retrieval treatment using Target retrieval Solution (Dako) by incubation at 80°C for 30 min and let to rest at room temperature for 30 min. After PBS washes, the sections were permeabilized 15 min in PBS-Triton 1% and then blocked for 1 hour at room temperature in 3% serum/0.3% PBS-Triton under gentle shaking. The sections were then incubated with the primary antibodies diluted (as described in supplementary table 1) in blocking solution overnight at 4°C. The next day, the sections were washed three times in PBS, followed by incubation in secondary antibody diluted in blocking solution for 2 hour at room temperature under gentle shaking. The sections were then washed in PBS, incubated in Hoechst 33342 solution (1:1000 in PBS, Molecular Probes/Life Technologies #H3570) for 10 min at room temperature. After a final PBS wash, the sections were mounted onto SuperFrost Plus adhesion slides (Epredia), let them to dry before adding Mounting medium (Prolong Gold Antifade Mounting medium, Invitrogen) and covered with a coverslip. Pictures were taken using a Zeiss LSM800-Airy confocal microscope.

For anti-DNMT3A and AIF1/IBA1 staining, the free-floating mice brain sections were washed several times with 1× Tris-buffered saline (TBS) to remove the cryoprotectant solution. For DNMT3A staining, an intermediate antigen retrieval step was required before blocking. To this end, the brain sections were incubated in sodium citrate solution (NaCi, 10 mM, pH 6.0) for 30 min at 80 ^°^C, allowed to recover for 10 min at room temperature, and then rinsed several times in TBS to remove the excess of NaCi.

Then the sections were subsequently incubated in a blocking solution containing 5 % donkey serum (Jackson ImmunoResearch Laboratories, West Grove, PA) and 0.1 % Triton X-100, for 1 hour at room temperature. Incubation with the primary antibodies ensued for 48 hours at 4 ^°^C. The sections were then rinsed with TBS and incubated with the appropriate secondary antibodies for 1 hour at room temperature. Hoechst 33342 was added for nuclear counterstaining. Finally, the sections were mounted onto microscope slides using ProLong Gold anti-fade reagent (Molecular probes/Life Technologies) as mounting media, coverslipped, and allowed to dry at room temperature. Image acquisition was carried out using a Zeiss LSM700 confocal laser scanning microscopy equipped with ZEN Zeiss software.

### Tumor volume calculation

Assessment of tumor size and microglial occupancy outcome were blindly analyzed by experimenter independent from the one who performed animal surgeries. Using a preestablish inclusion criteria, animals presenting tumor size below 1 mm^3^ were excluded from the analysis. Volumes in mm^3^ were calculated in coronal sections using the Zeiss software from the GFP-positive and Iba1-positive areas according to the Cavalieri principle using the following algorithm: *V* = Σ*A* × *P* × *T*, where *V* = total volume, Σ*A* = the sum of area measurements, *P* = the inverse of the sampling fraction, and *T* = the section thickness.

### Statistical analyses

All statistical analyses were conducted using Prism 8 (v.9.3.1, GraphPad Software). Results were tested for statistical significance using one-way ANOVA and Bonferroni’s test to correct for multiple comparisons. If two conditions were to be compared, two-tailed paired or unpaired Student’s *t* test was used. Results are presented as the mean ± s.e.m. Differences were considered statistically significant if *P* values were less than 0.05. Raw data are available as Source Data.

### Data availability

RNA-Seq and ChIP-Seq data will be deposited in the Gene Expression Omnibus prior to article publication. All other data are available from the corresponding author upon reasonable request.

## Acknowledgments

We thank G. Brown (University of Cambridge) for the BV2 cell line; O. Hermanson (Karolinska Institutet) for the C6 cell line; M. Nister (Karolinska Institutet) for the U-87MG cell line; M. Schultzberg (Karolinska Institutet) for the HMC3 cell line. The authors would like to thank the core facility at Novum, BEA, Bioinformatics and Expression Analysis, which is supported by the board of research at the Karolinska Institute and the research committee at the Karolinska hospital. This research is supported by the Swedish Research Council and the Swedish Brain Foundation, the Cancer Research Funds of Radiumhemmet, the Strategic Research Programme in Cancer (StratCan), the Strategic Research Programme in Neuroscience (StratNeuro), (B.J.), the Swedish Cancer Society, the Swedish Childhood Cancer Foundation (K.B., and B.J.), the Karolinska Institutet Foundation (G.V.C, L.F., and B.J.), the Åke Wibergs Stiftelse, the Hedlunds Foundation (M.C.), the Swedish governmental grants for researchers working in healthcare (K.B.).

## Authors contributions

M.C. designed and performed the experiments and analyzed the data except otherwise indicated. A.N.M., A.F., C.R., A.M.O., L.M.C. and K.B. performed the mice experiments. M.C., A.N.M, C.R. and A.M.O. performed mice brain preparation, staining and confocal analysis. G.V.C. participated in the confocal analysis. S.K., Y.L. and P.U. performed the RNA-scope experiment and associated confocal analysis. A.D. performed the data analysis of the RNA-seq and ChIP seq. M.S., L.F., and B.J. contributed to bioinformatic analysis of transcriptomic datasets. C.H. and F.K. generated and provided the ASOs and performed the ASOs mice tolerability data. M.C. and B.J. initiated, conceptualized, and designed the study, analyzed, and interpreted the data and wrote the first draft of the paper. All authors discussed the results and commented or edited the paper.

---

**Supplementary Table 1.**
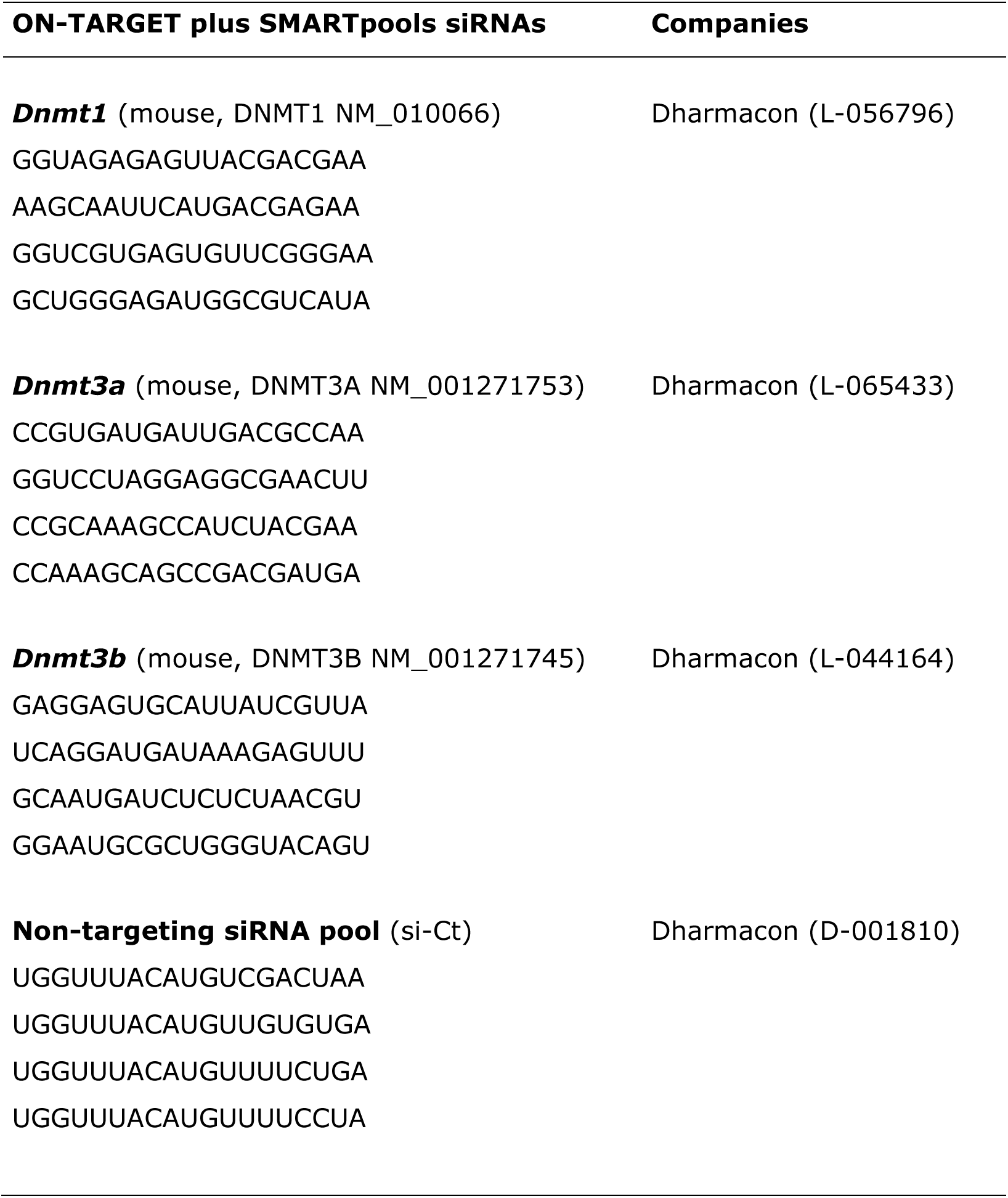
ON-TARGETplus SMART pool small interfering RNAs used in this study.

**Supplementary Table 2.**
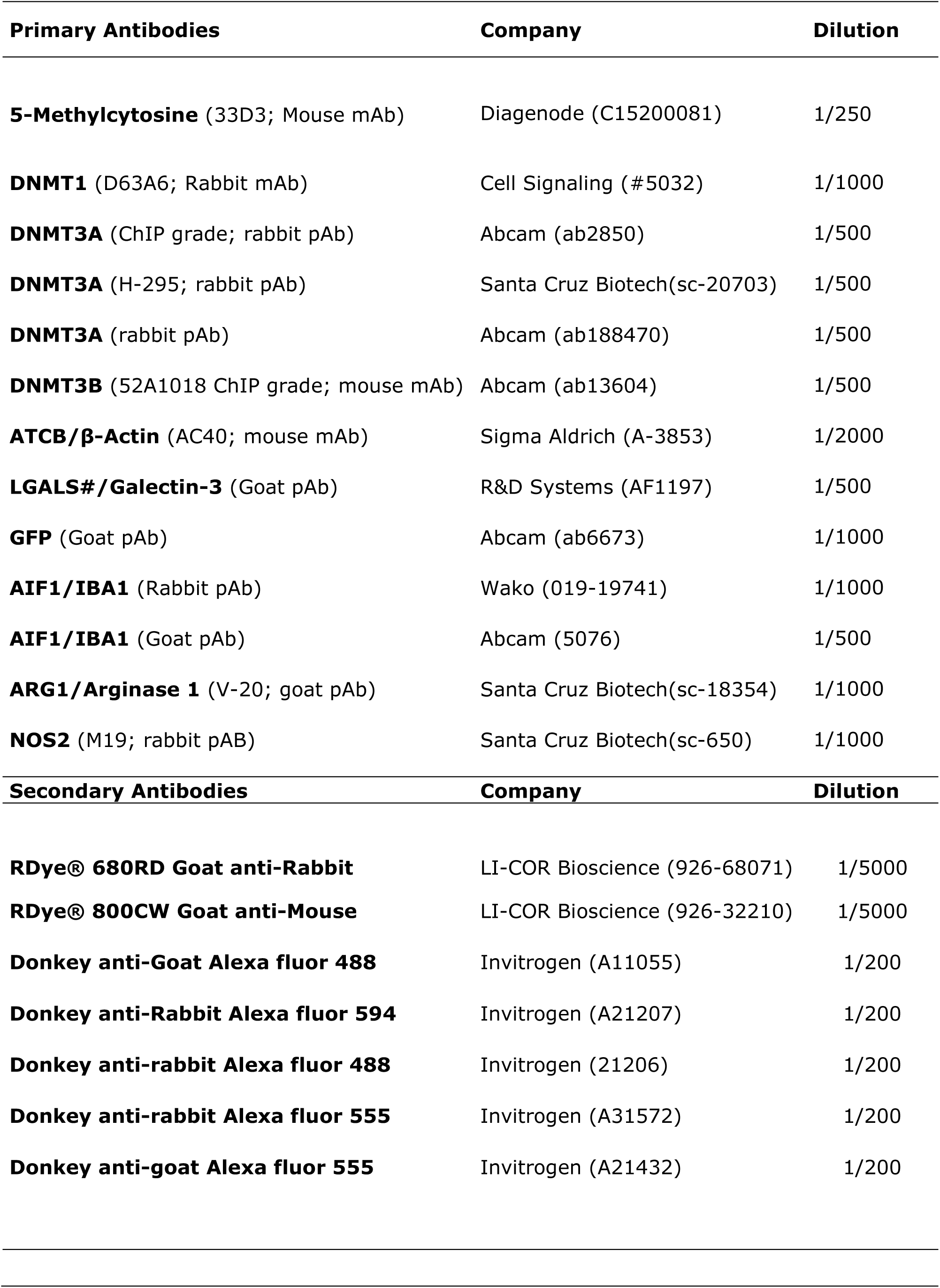
Antibodies used in this study.

**Supplementary Table 3.**
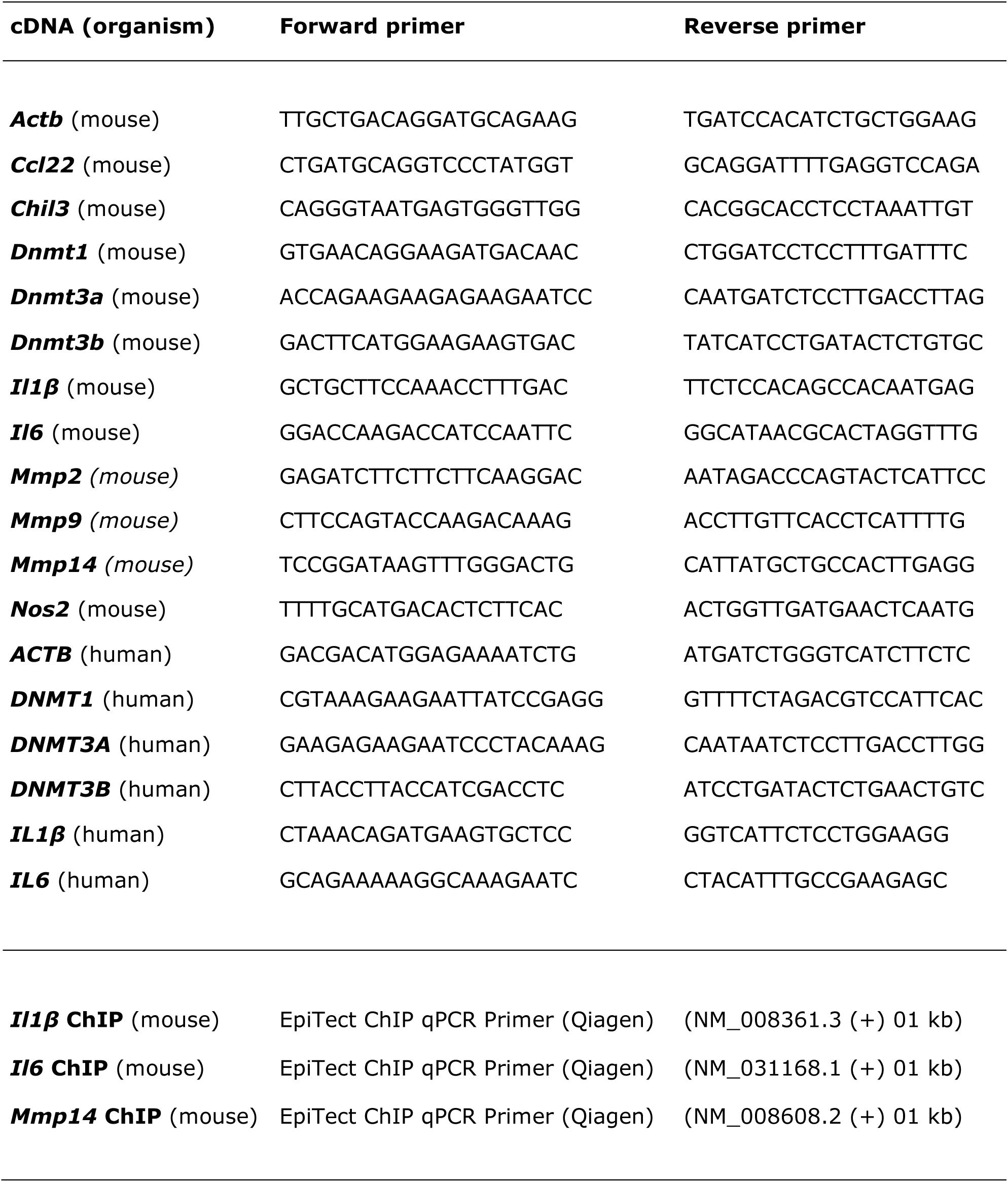
Primer sequences.

## Extended Data Figure legends

**Extended Data Figure 1 | Transcriptomic response of BV2 microglia exposed to C6 glioma cells in coculture for 2 or 4 hours.**

**a**,**b**, Volcano plot representations of the RNA-seq data comparing gene expression in BV2 microglia cocultured with C6 glioma cells versus monoculture at 2h (**a**) and 4h (**b**) time points. Blue dots represent less expressed genes, whereas red dots represent genes that are higher expressed in C6 glioma cells-stimulated BV2 microglia, respectively. **c**,**d**, Cytoscape network analysis of enriched GO term clusters generated from upregulated genes in C6 glioma cells-stimulated BV2 microglia at the 2h (**c**) and 4h (**d**) time points with significant FDR (< 0.05). Nodes represent GO terms, clusters are nodes grouped based on similarity. Node size corresponds to the number of genes. Node color corresponds to the significance of correlation, where the darker the color is, the smaller the FDR values gets. Lines represent the number of genes overlapping between nodes.

**Extended Data Figure 2 | Decreased DNMT3A expression, and induction of *IL1β* expression are also observed in a human HMC3 microglia-U87-MG glioblastoma coculture setup**

**a**, RTqPCR analysis of *IL1b* and *IL6* mRNA expression levels in human HMC3 microglia exposed to coculture with U87-MG glioblastoma cells for 2h, 4h, and 6h (n=3; Data are shown as mean ± sem). **b**,**c**, Immunoblot analysis (**b**), with quantifications (**c**), of DNMT3 and β-actin (ACTB) expression levels in HMC3 microglia in monoculture or in coculture with U87-MG glioblastoma cells at the indicated time-points (n=3; Data are shown as mean ± sem). **d**, RTqPCR analysis of *DNMT1*, *DNMT3A*, and *DNMT3B* mRNA expression levels in human HMC3 microglia exposed to coculture with U87-MG glioblastoma cells for 2h, 4h, and 6h (n=3; Data are shown as mean ± sem).

**Extended Data Figure 3 | Stimulation of BV2 microglia with LPS or IL4 treatments do not mimic the effects observed upon exposure to C6 glioma cells.**

**a**,**b**, Immunoblot analysis (**a**) of DNMT3, arginase-1 (ARG1), nitric oxide synthase 2 (NOS2), and β-actin (ACTB) expression levels in BV2 microglia treated with LPS (100ng/ml) or IL4 (20ng/ml) for 2h, 4h, 6h, and 24 h with quantifications for DNMT3A (**b**) (n=3; Data are shown as mean ± sem). **c**,**d**,**e**,**f**, RTqPCR analysis of mRNA expression levels in BV2 microglia treated with LPS (**c,e**) or IL4 (**d,f**) for 2h, 4h, 6h, and 24 h of genes frequently associated to a microglial tumor-supportive phenotype, *i.e*., *Ccl22* and *Chil3/Ym1* and proinflammatory cytokines, *i.e*., *Il1b* and *Il6* (**c,d**) or *Dnmt1*, *Dnmt3a*, and *Dnmt3b* (**e**,**f**). (n=3; Data are shown as mean ± sem).

**Extended Data Figure 4 | Repression of *Dnmt1* or *Dnmt3b* expression in BV2 microglia do not mimic the effect observed upon *Dnmt3a* silencing.**

**a,** RTqPCR analysis of *Dnmt1*, *Dnmt3a*, *Dnmt3b* mRNA expression levels in BV2 microglia transfected with a pool of siRNAs targeting *Dnmt3a* expression (siRNA-*Dnmt3a*), as compared to cells transfected with a pool of non-targeting siRNAs control (siRNA-Control/Ctrl, negative control), set as 1. (n=3; Data are shown as mean ± sem). **b,** RTqPCR analysis of *Dnmt1*, *Dnmt3a*, *Dnmt3b* mRNA expression levels in BV2 microglia transfected with a pool of siRNAs targeting *Dnmt1* expression (siRNA-*Dnmt1*), or *Dnmt3b* expression (siRNA-*Dnmt3b*), as compared to cells transfected with siRNA-Control/Ctrl, negative control, set as 1. (n=3; Data are shown as mean ± sem). **c**,**d**, Immunoblot analysis of DNMT1 (**c**), or DNMT3B (**d**) together with β-actin (ACTB) expression levels in mock transfected (-) BV2 microglia, or cells transfected with siRNA-Ctrl, siRNA-*Dnmt1* (**c**) or siRNA-*Dnmt3b* (**d**) (n=3; Data are shown as mean ± sem). **e**, Analysis of cell migration capability of mock transfected (-) BV2 microglia, or cells transfected with siRNA-Ctrl, siRNA-*Dnmt1,* or siRNA-*Dnmt3b.* **f**, Analysis of C6 glioma cell migration capability toward mock transfected (-) BV2 microglia, or cells transfected with siRNA-Ctrl, siRNA-*Dnmt1,* or siRNA-*Dnmt3b* (n=3; Data are shown as mean ± sem).

**Extended Data Figure 5 | *In vivo*, co-injection of shRNA *Dnmt3a* BV2 microglia with GL261 glioblastoma cells reduces tumor growth.**

**a,b,** Immunoblot analysis (**a**), with quantifications (**b**), of DNMT3A and β-actin (ACTB) expression levels in BV2 cell lines generated to express constitutively a vector encoding for shRNA targeting *Dnmt3a* expression (2 different lentiviral clones #1 and #2 are depicted) or an empty vector used as control (Ct). **c**, Confocal immunofluorescence imaging showing GFP-GL261 glioblastoma cells (green), and IBA1-expressing microglia (red) in mouse brain tissues 2 weeks after the intracranial injection GL261 cancer cells alone, or a mix with shRNA *Dnmt3a* BV2 microglia. Hoechst was used for nuclear counterstain (blue). **d**, Quantification of the tumor sizes in the above-described conditions. (n=8 mice per group. Data are shown as box plot).

**Extended Data Figure 6 | *In vivo*, antisense oligonucleotides targeting *Dnmt3a* efficiently reduces *Dnmt3a* expression and promote microglial galectin3 expression.**

**a, b,** RTqPCR analysis of *Dnmt3a* mRNA expression level in the cortex (left panel) or spinal cord (right panel) of mice injected with PBS (used as control) or with ASOs targeting *Dnmt3a* (ASO-1 and ASO-2) and sacrificed at 2 weeks (**a**) or 8 weeks (**b**) after injection, ratio to vehicle, average of two independent experiments. **c**, Confocal immunofluorescence imaging showing galectin-3 (LGALS3, green), and IBA1-expressing microglia (red) in brain tissues of mice injected with ASO-Ct or ASO targeting DNMT3A (ASO-1 and ASO-2). Hoechst (blue) as nuclear counterstain. (n=3, Scale bars 1000µm). Zooms in merge images corresponding to the white doted boxes are presented in the right hand side. **d**, Confocal immunofluorescence imaging of tumors formed 2 weeks post injection of GFP-GL261 glioma cells (green) in mice previously treated with PBS, or ASO-Ct (as illustrated in c) together with an immunostaining for the microglial marker IBA1 (red) and Hoechst used as nuclear counterstain (blue). **e**, Quantification of 2 weeks old GL261 tumor volume in mice injected with PBS or ASO-Ct, 2 weeks prior injection of GFP-GL261 cancer cells. (n=9/10 mice per group; Data are presented as box plot)(Scale bars 100µm).

