## supplementary figures for "Glioma-induced DNMT3A-dependent reduction of DNA methylation in microglia promotes a transient anti-tumoral phenotype"

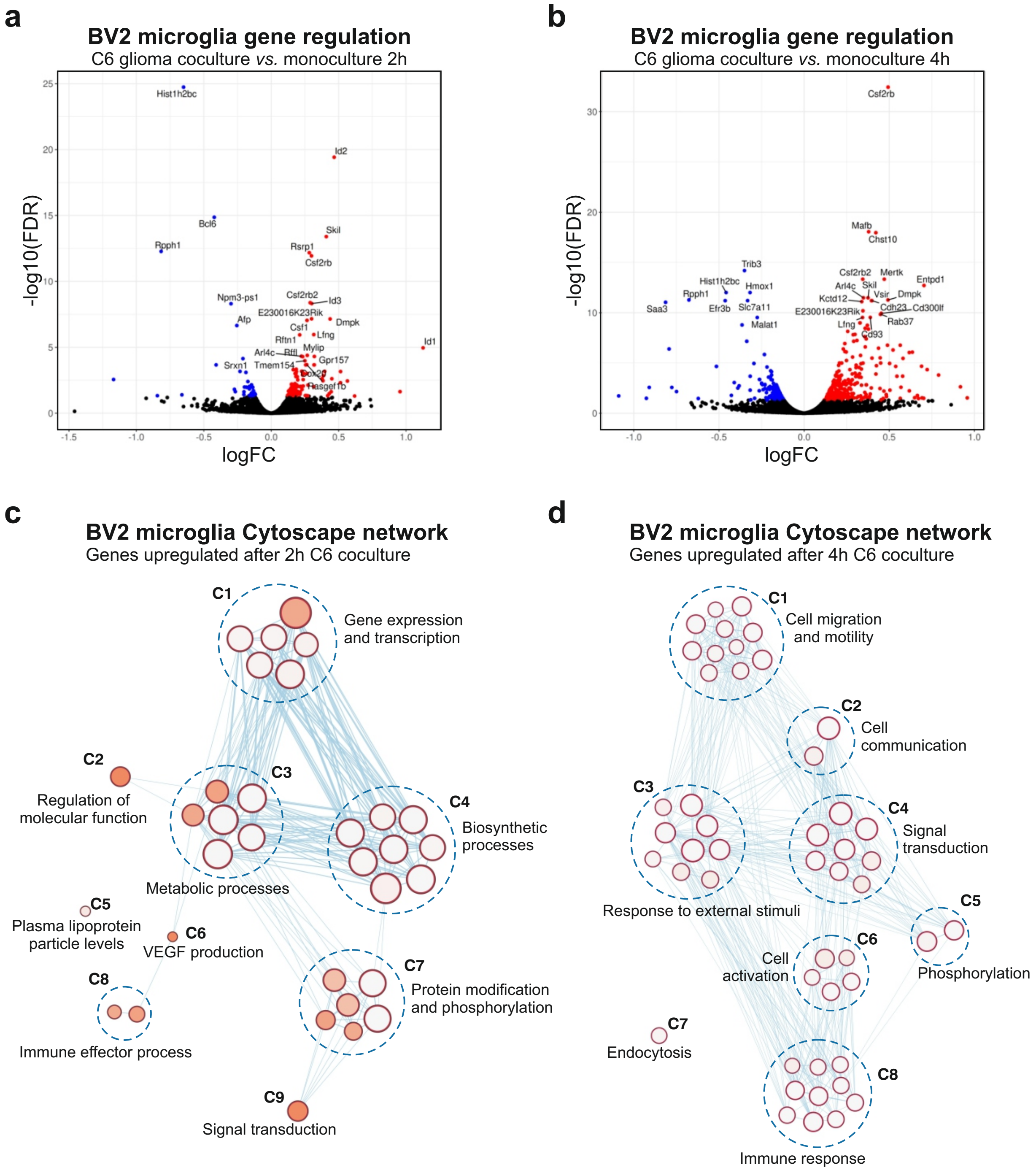


**Supplementary Figure 1. Cheray et al.**

**Supplementary Figure 1 | Transcriptomic response of BV2 microglia exposed to C6 glioma cells in coculture for 2 or 4 hours.**

**a**,**b**, Volcano plot representations of the RNA-seq data comparing gene expression in BV2 microglia cocultured with C6 glioma cells versus monoculture at 2h (**a**) and 4h (**b**) time points. Blue dots represent less expressed genes, whereas red dots represent genes that are higher expressed in C6 glioma cells-stimulated BV2 microglia, respectively. **c**,**d**, Cytoscape network analysis of enriched GO term clusters generated from upregulated genes in C6 glioma cells-stimulated BV2 microglia at the 2h (**c**) and 4h (**d**) time points with significant FDR (< 0.05). Nodes represent GO terms, clusters are nodes grouped based on similarity. Node size corresponds to the number of genes. Node color corresponds to the significance of correlation, where the darker the color is, the smaller the FDR values gets. Lines represent the number of genes overlapping between nodes.

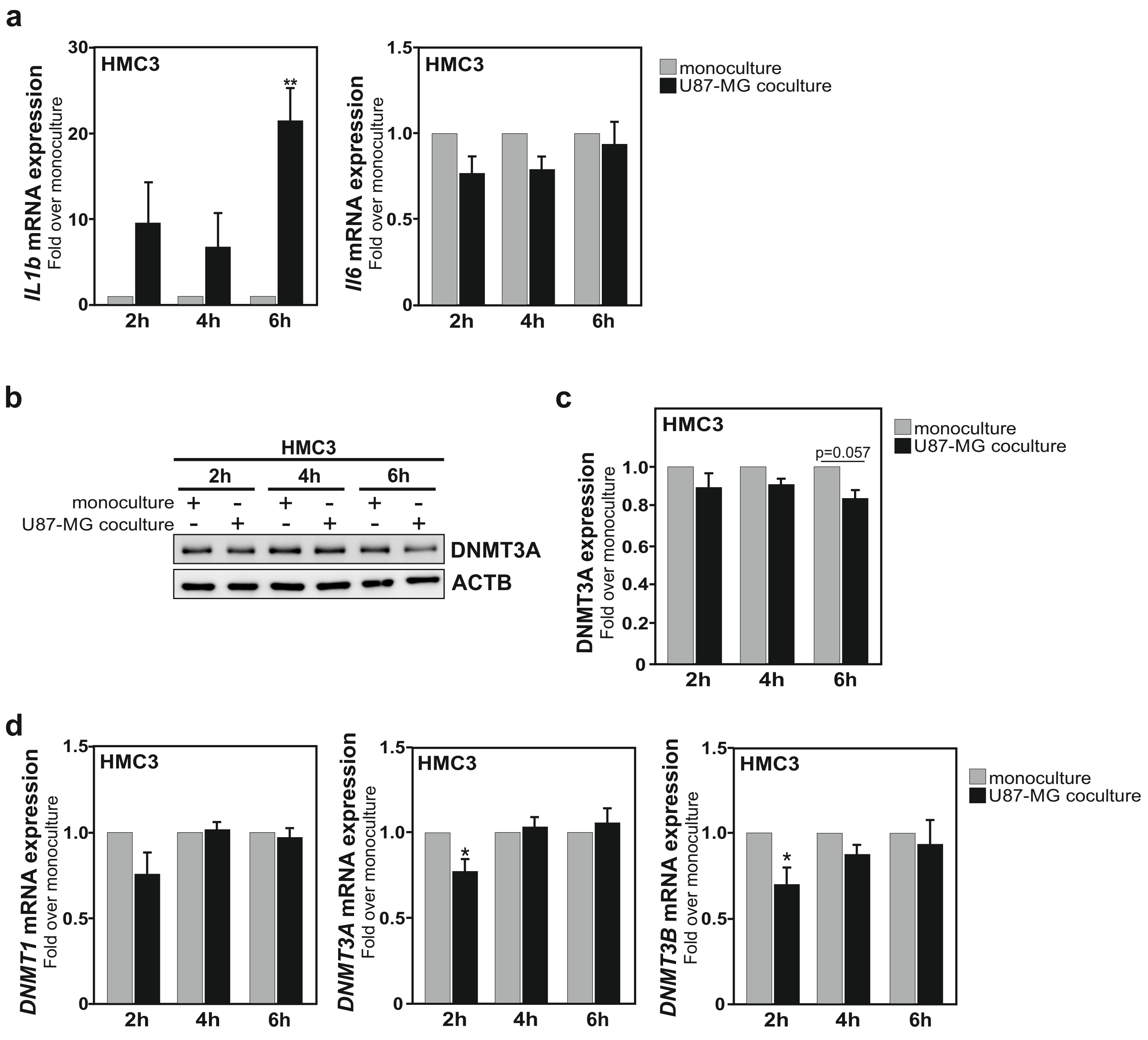


**Supplementary Figure 2. Cheray et al.**

**Supplementary Figure 2 | Decreased DNMT3A expression, and induction of *IL1β* expression are also observed in a human HMC3 microglia-U87-MG glioblastoma coculture setup**

**a**, RTqPCR analysis of *IL1b* and *IL6* mRNA expression levels in human HMC3 microglia exposed to coculture with U87-MG glioblastoma cells for 2h, 4h, and 6h (n=3; Data are shown as mean ± sem). **b**,**c**, Immunoblot analysis (**b**), with quantifications (**c**), of DNMT3 and β-actin (ACTB) expression levels in HMC3 microglia in monoculture or in coculture with U87-MG glioblastoma cells at the indicated time-points (n=3; Data are shown as mean ± sem). **d**, RTqPCR analysis of *DNMT1*, *DNMT3A*, and *DNMT3B* mRNA expression levels in human HMC3 microglia exposed to coculture with U87-MG glioblastoma cells for 2h, 4h, and 6h (n=3; Data are shown as mean ± sem).

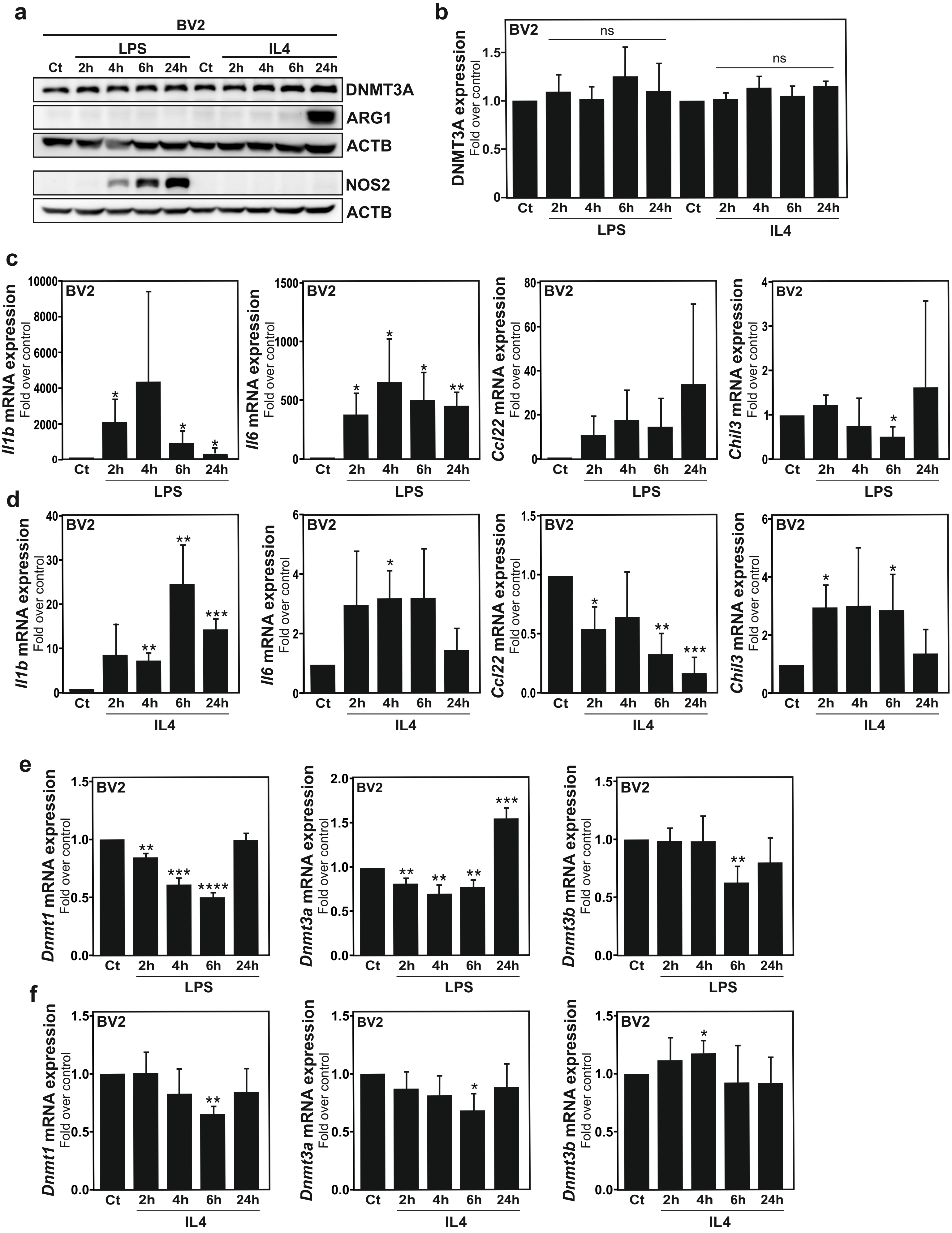


Supplementary Figure 3. Cheray et al.

**Supplementary Figure 3 | Stimulation of BV2 microglia with LPS or IL4 treatments do not mimic the effects observed upon exposure to C6 glioma cells.**

**a**,**b**, Immunoblot analysis (**a**) of DNMT3, arginase-1 (ARG1), nitric oxide synthase 2 (NOS2), and β-actin (ACTB) expression levels in BV2 microglia treated with LPS or IL4 for 2h, 4h, 6h, and 24 h with quantifications for DNMT3A (**b**) (n=3; Data are shown as mean ± sem). **c**,**d**,**e**,**f**, RTqPCR analysis of mRNA expression levels in BV2 microglia treated with LPS (**c**,**e**) or IL4 (**d**,**f**) for 2h, 4h, 6h, and 24 h of genes frequently associated to a microglial tumor-supportive phenotype, *i.e.*, *Ccl22* and *Chil3/Ym1* and proinflammatory cytokines, *i.e.*, *Il1b* and *Il6* (**c**,**d**) or *Dnmt1*, *Dnmt3a*, and *Dnmt3b* (**e**,**f**). (n=3; Data are shown as mean ± sem).

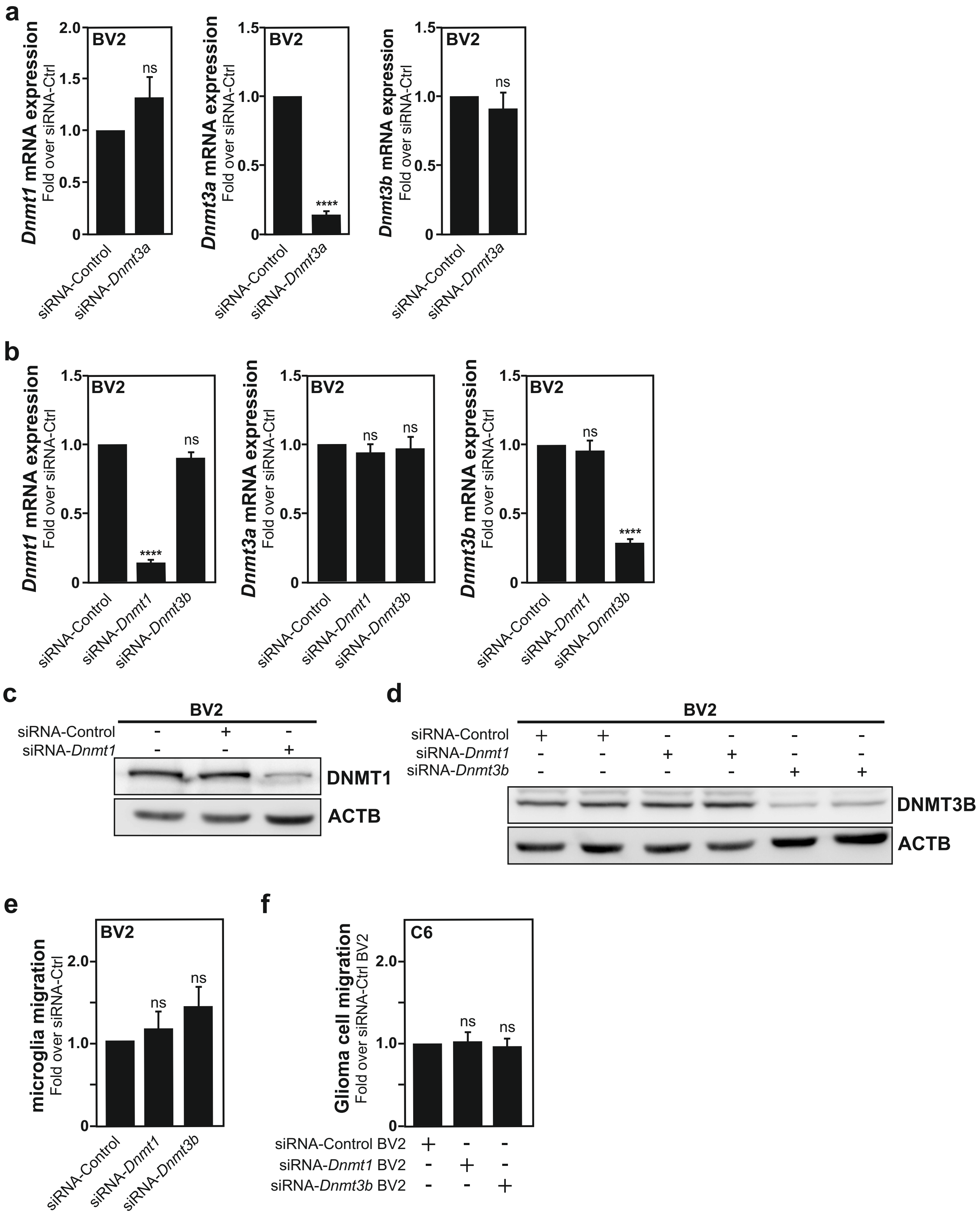


**Supplementary Figure 4. Cheray et al.**

**Supplementary Figure 4 | Repression of *Dnmt1* or *Dnmt3b* expression in BV2 microglia do not mimic the effect observed upon *Dnmt3a* silencing.**

**a,** RTqPCR analysis of *Dnmt1*, *Dnmt3a*, *Dnmt3b* mRNA expression levels in BV2 microglia transfected with a pool of siRNAs targeting *Dnmt3a* expression (siRNA-*Dnmt3a*), as compared to cells transfected with a pool of non-targeting siRNAs control (siRNA-Control/Ctrl, negative control), set as 1. (n=3; Data are shown as mean ± sem). **b,** RTqPCR analysis of *Dnmt1*, *Dnmt3a*, *Dnmt3b* mRNA expression levels in BV2 microglia transfected with a pool of siRNAs targeting *Dnmt1* expression (siRNA-*Dnmt1*), or *Dnmt3b* expression (siRNA-*Dnmt3b*), as compared to cells transfected with siRNA-Control/Ctrl, negative control, set as 1. (n=3; Data are shown as mean ± sem). **c**,**d**, Immunoblot analysis of DNMT1 (**c**), or DNMT3B (**d**) together with β-actin (ACTB) expression levels in mock transfected (-) BV2 microglia, or cells transfected with siRNA-Ctrl, siRNA-*Dnmt1* (**c**) or siRNA-*Dnmt3b* (**d**) (n=3; Data are shown as mean ± sem). **e**, Analysis of cell migration capability of mock transfected (-) BV2 microglia, or cells transfected with siRNA-Ctrl, siRNA-*Dnmt1,* or siRNA-*Dnmt3b.* **f**, Analysis of C6 glioma cell migration capability toward mock transfected (-) BV2 microglia, or cells transfected with siRNA-Ctrl, siRNA-*Dnmt1,* or siRNA-*Dnmt3b* (n=3; Data are shown as mean ± sem).

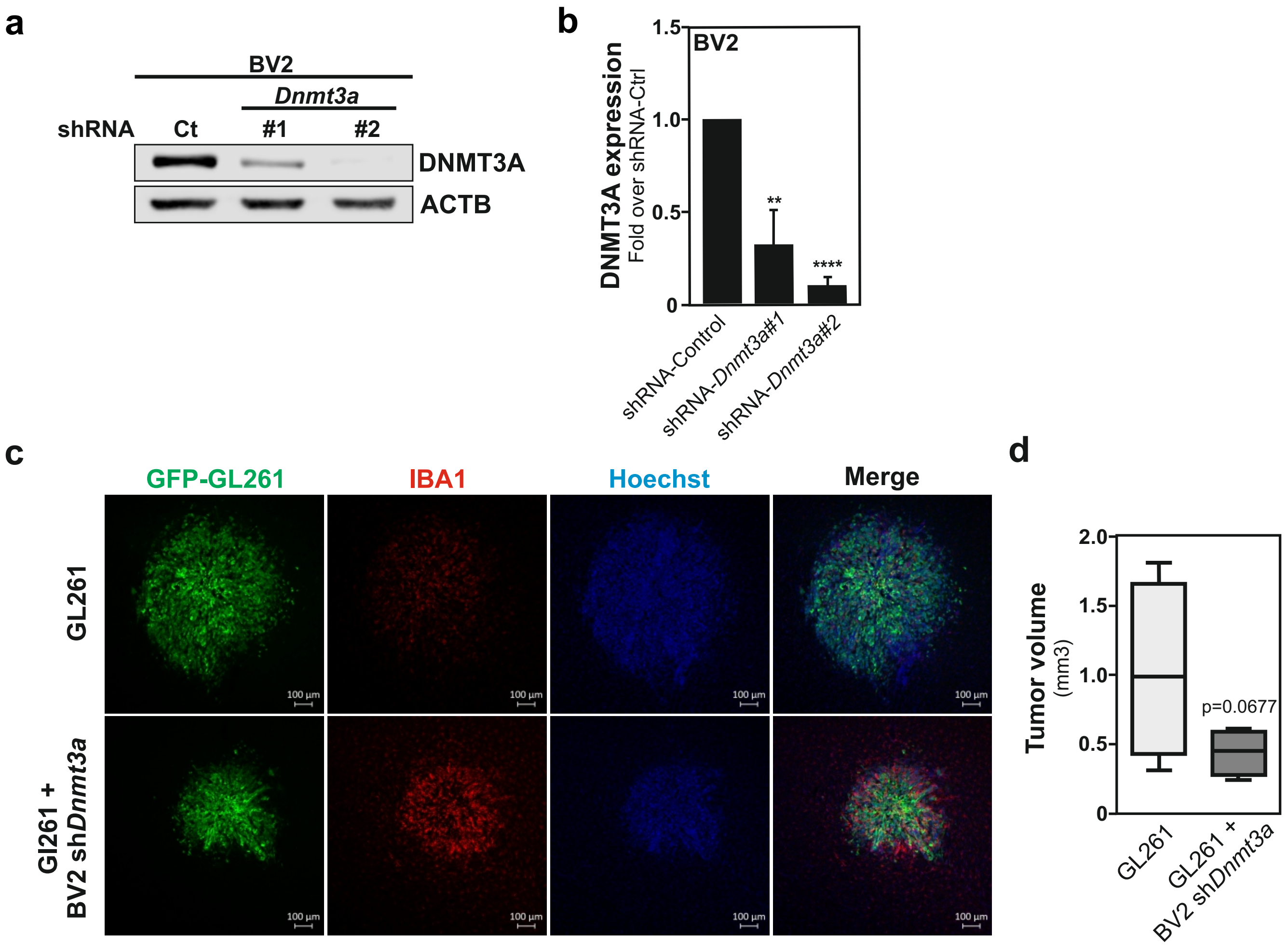


**Supplementary Figure 5. Cheray et al.**

**Supplementary Figure 5 | *In vivo*, co-injection of shRNA *Dnmt3a* BV2 microglia with GL261 glioblastoma cells reduces tumor growth.**

**a,b,** Immunoblot analysis (**a**), with quantifications (**b**), of DNMT3A and β-actin (ACTB) expression levels in BV2 cell lines generated to express constitutively a vector encoding for shRNA targeting *Dnmt3a* expression (2 different lentiviral clones #1 and #2 are depicted) or an empty vector used as control (Ct). **c**, Confocal immunofluorescence imaging showing GFP-GL261 glioblastoma cells (green), and IBA1-expressing microglia (red) in mouse brain tissues 2 weeks after the intracranial injection GL261 cancer cells alone, or a mix with shRNA *Dnmt3a* BV2 microglia. Hoechst was used for nuclear counterstain (blue). **d**, Quantification of the tumor sizes in the above-described conditions. (n=8 mice per group. Data are shown as box plot).

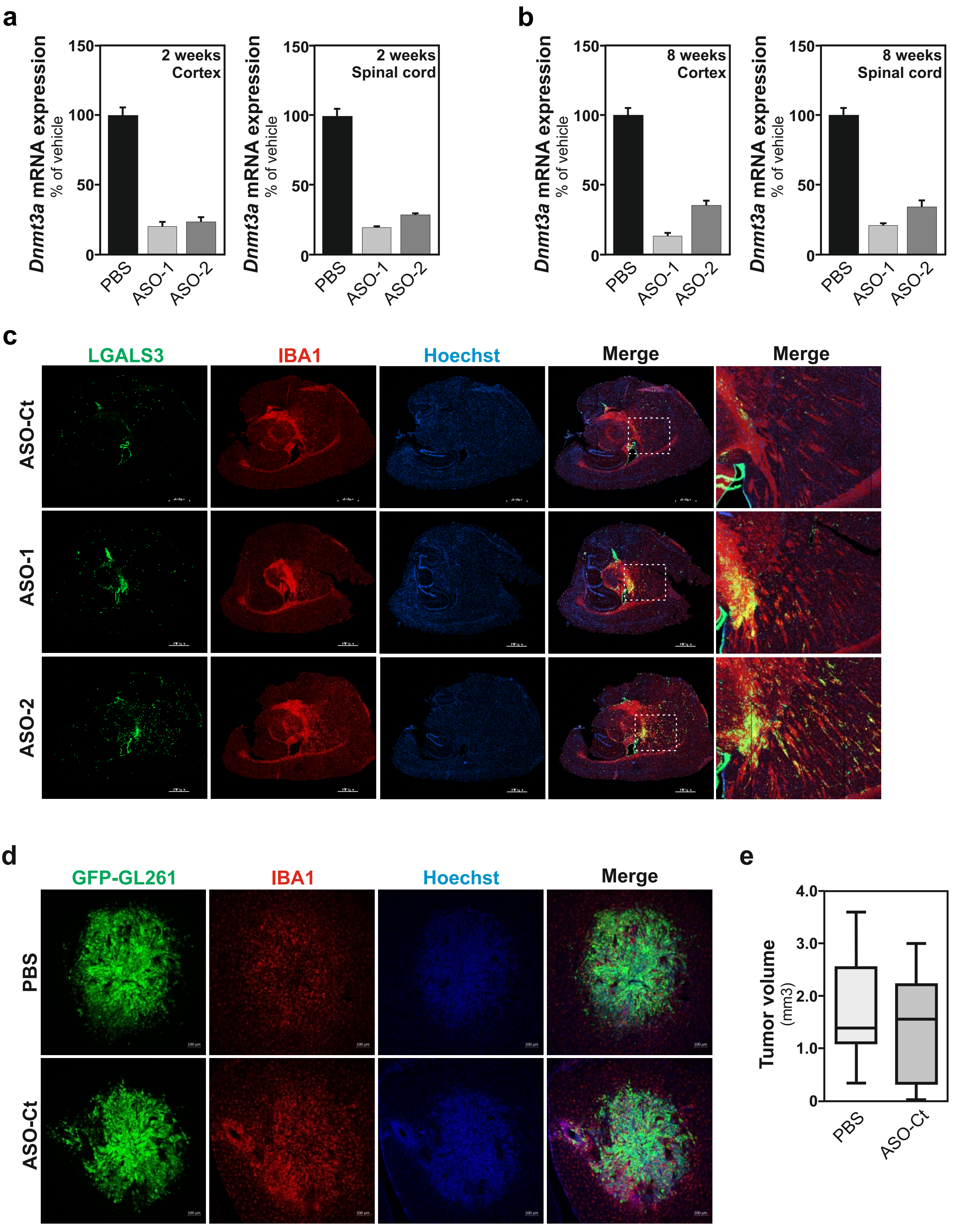


**Supplementary Figure 6. Cheray et al.**

**Supplementary Figure 6 | *In vivo*, antisense oligonucleotides targeting *Dnmt3a* efficiently reduces *Dnmt3a* expression and promote microglial galectin3 expression.**

**a, b,** RTqPCR analysis of *Dnmt3a* mRNA expression level in the cortex (left panel) or spinal cord (right panel) of mice injected with PBS (used as control) or with ASOs targeting *Dnmt3a* (ASO-1 and ASO-2) and sacrificed at 2 weeks (**a**) or 8 weeks (**b**) after injection, ratio to vehicle, average of two independent experiments. **c**, Confocal immunofluorescence imaging showing galectin-3 (LGALS3, green), and IBA1-expressing microglia (red) in brain tissues of mice injected with ASO-Ct or ASO targeting DNMT3A (ASO-1 and ASO-2). Hoechst (blue) as nuclear counterstain. (n=3, Scale bars 1000µm). Zooms in merge images corresponding to the white doted boxes are presented in the right hand side. **d**, Confocal immunofluorescence imaging of tumors formed 2 weeks post injection of GFP-GL261 glioma cells (green) in mice previously treated with PBS, or ASO-Ct (as illustrated in c) together with an immunostaining for the microglial marker IBA1 (red) and Hoechst used as nuclear counterstain (blue). **e**, Quantification of 2 weeks old GL261 tumor volume in mice injected with PBS or ASO-Ct, 2 weeks prior injection of GFP-GL261 cancer cells. (n=9/10 mice per group; Data are presented as box plot)(Scale bars 100µm).
