## supplementary tables for "Glioma-induced DNMT3A-dependent reduction of DNA methylation in microglia promotes a transient anti-tumoral phenotype"

**Supplementary Table 1 |ON-TARGETplus SMART pool small interfering RNAs used in this study.**

| ON-TARGET plus SMARTpools siRNAs | Companies |
|---|---|
| ***Dnmt1*** (mouse, DNMT1 NM_010066) | Dharmacon (L-056796) |
| GGUAGAGAGUUACGACGAA | |
| AAGCAAUUCAUGACGAGAA | |
| GGUCGUGAGUGUUCGGGAA | |
| GCUGGGAGAUGGCGUCAUA | |
| ***Dnmt3a*** (mouse, DNMT3A NM_001271753) | Dharmacon (L-065433) |
| CCGUGAUGAUUGACGCCAA | |
| GGUCCUAGGAGGCGAACUU | |
| CCGCAAAGCCAUCUACGAA | |
| CCAAAGCAGCCGACGAUGA | |
| ***Dnmt3b*** (mouse, DNMT3B NM_001271745) | Dharmacon (L-044164) |
| GAGGAGUGCAUUAUCGUUA | |
| UCAGGAUGAUAAAGAGUUU | |
| GCAAUGAUCUCUCUAACGU | |
| GGAAUGCGCUGGGUACAGU | |
| **Non-targeting siRNA pool** (si-Ct) | Dharmacon (D-001810) |
| UGGUUUACAUGUCGACUAA | |
| UGGUUUACAUGUUGUGUGA | |
| UGGUUUACAUGUUUUCUGA | |
| UGGUUUACAUGUUUUCCUA | |

**Supplementary Table 2 | Antibodies used in this study**

| **Primary Antibodies** | **Company** | **Dilution** |
|---|---|---|
| **5-Methylcytosine** (33D3; Mouse mAb) | Diagenode (C15200081) | 1/250 |
| **DNMT1** (D63A6; Rabbit mAb) | Cell Signaling (#5032) | 1/1000 |
| **DNMT3A** (ChIP grade; rabbit pAb) | Abcam (ab2850) | 1/500 |
| **DNMT3A** (H-295; rabbit pAb) | Santa Cruz Biotech(sc-20703) | 1/500 |
| **DNMT3A** (rabbit pAb) | Abcam (ab188470) | 1/500 |
| **DNMT3B** (52A1018 ChIP grade; mouse mAb) | Abcam (ab13604) | 1/500 |
| **ATCB/β-Actin** (AC40; mouse mAb) | Sigma Aldrich (A-3853) | 1/2000 |
| **LGALS#/Galectin-3** (Goat pAb) | R&D Systems (AF1197) | 1/500 |
| **GFP** (Goat pAb) | Abcam (ab6673) | 1/1000 |
| **AIF1/IBA1** (Rabbit pAb) | Wako (019-19741) | 1/1000 |
| **AIF1/IBA1** (Goat pAb) | Abcam (5076) | 1/500 |
| **ARG1/Arginase 1** (V-20; goat pAb) | Santa Cruz Biotech(sc-18354) | 1/1000 |
| **NOS2** (M19; rabbit pAB) | Santa Cruz Biotech(sc-650) | 1/1000 |

| **Secondary Antibodies** | **Company** | **Dilution** |
|---|---|---|
| **RDye® 680RD Goat anti-Rabbit** | LI-COR Bioscience (926-68071) | 1/5000 |
| **RDye® 800CW Goat anti-Mouse** | LI-COR Bioscience (926-32210) | 1/5000 |
| **Donkey anti-Goat Alexa fluor 488** | Invitrogen (A11055) | 1/200 |
| **Donkey anti-Rabbit Alexa fluor 594** | Invitrogen (A21207) | 1/200 |
| **Donkey anti-rabbit Alexa fluor 488** | Invitrogen (21206) | 1/200 |
| **Donkey anti-rabbit Alexa fluor 555** | Invitrogen (A31572) | 1/200 |
| **Donkey anti-goat Alexa fluor 555** | Invitrogen (A21432) | 1/200 |

**Supplementary Table 3 | Primer sequences**

All sequences are given 5´to 3´

| cDNA (organism) | Forward primer | Reverse primer |
|---|---|---|
| ***Actb*** (mouse) | TTGCTGACAGGATGCAGAAG | TGATCCACATCTGCTGGAAG |
| ***Ccl22*** (mouse) | CTGATGCAGGTCCCTATGGT | GCAGGATTTTGAGGTCCAGA |
| ***Chil3*** (mouse) | CAGGGTAATGAGTGGGTTGG | CACGGCACCTCCTAAATTGT |
| ***Dnmt1*** (mouse) | GTGAACAGGAAGATGACAAC | CTGGATCCTCCTTTGATTTC |
| ***Dnmt3a*** (mouse) | ACCAGAAGAAGAGAAGAATCC | CAATGATCTCCTTGACCTTAG |
| ***Dnmt3b*** (mouse) | GACTTCATGGAAGAAGTGAC | TATCATCCTGATACTCTGTGC |
| ***Il1β*** (mouse) | GCTGCTTCCAAACCTTTGAC | TTCTCCACAGCCACAATGAG |
| ***Il6*** (mouse) | GGACCAAGACCATCCAATTC | GGCATAACGCACTAGGTTTG |
| ***Mmp2*** *(mouse)* | GAGATCTTCTTCTTCAAGGAC | AATAGACCCAGTACTCATTCC |
| ***Mmp9*** *(mouse)* | CTTCCAGTACCAAGACAAAG | ACCTTGTTCACCTCATTTTG |
| ***Mmp14*** *(mouse)* | TCCGGATAAGTTTGGGACTG | CATTATGCTGCCACTTGAGG |
| ***Nos2*** (mouse) | TTTTGCATGACACTCTTCAC | ACTGGTTGATGAACTCAATG |
| ***ACTB*** (human) | GACGACATGGAGAAAATCTG | ATGATCTGGGTCATCTTCTC |
| ***DNMT1*** (human) | CGTAAAGAAGAATTATCCGAGG | GTTTTCTAGACGTCCATTCAC |
| ***DNMT3A*** (human) | GAAGAGAAGAATCCCTACAAAG | CAATAATCTCCTTGACCTTGG |
| ***DNMT3B*** (human) | CTTACCTTACCATCGACCTC | ATCCTGATACTCTGAACTGTC |
| ***IL1β*** (human) | CTAAACAGATGAAGTGCTCC | GGTCATTCTCCTGGAAGG |
| ***IL6*** (human) | GCAGAAAAAGGCAAAGAATC | CTACATTTGCCGAAGAGC |
| ***Il1β* ChIP** (mouse) | EpiTect ChIP qPCR Primer (Qiagen) | (NM_008361.3 (+) 01 kb) |
| ***Il6* ChIP** (mouse) | EpiTect ChIP qPCR Primer (Qiagen) | (NM_031168.1 (+) 01 kb) |
| ***Mmp14* ChIP** (mouse) | EpiTect ChIP qPCR Primer (Qiagen) | (NM_008608.2 (+) 01 kb) |
